# Mechanistic Understanding of Protein–MOF Integration through Surfactant-Driven Interfacial Design

**DOI:** 10.1101/2025.11.11.687871

**Authors:** Ehsan Rashidniyaghi, Mohammad Khavani, Carlie Coerver, Ruibin Liang, Raheleh Ravanfar

## Abstract

Integration of proteins into metal–organic frameworks (Protein@MOF) represents an effective method for protein stabilization, with rising demand across material and biomedical sciences. However, the molecular mechanism of protein–MOF interactions remains unsettled due to challenges in developing a general platform to systematically investigate such interactions, hindering improvements in their chemical and physical properties. Here, we develop a surfactant-guided strategy to modulate the assembly of protein@MOF through interfacial design. We discovered that the interfacial environment between proteins and MOFs is the primary factor determining encapsulation efficiency, structural retention, and functional performance. Lipid-based non-ionic surfactants such as glycerol monooleate (GMO) increase the protein’s solvent-accessible surface area (SASA), suggesting partial remodeling of the protein surface and hydration shell. GMO at the interface of protein@MOF results in a 20% improvement in protein encapsulation and a 30% increase in MOF growth rate. All-atom molecular dynamics simulations reveal domain-specific interactions between GMO and flexible surface residues on protein in a concentration-dependent manner, involving both electrostatic and hydrophobic contacts. This work offers new molecular insights into how surfactant-driven interfacial design fine-tunes the stability of protein@MOF, laying the foundation for robust alternatives to lipid nanodiscs for membrane protein stabilization, and protein-based platforms for drug-delivery, biocatalysis, and biosensing.

## 1. Introduction

Proteins drive essential biological processes in living systems with exceptional efficiency, selectivity, and structural adaptability.^[1]^ These traits have motivated widespread efforts to harness proteins as functional materials in applications ranging from catalysis and sensing to targeted therapeutics.^[2]^ In nature, protein-based materials, spanning from cytoskeletal filaments to silk fibers, exemplify how hierarchical structure and dynamic responsiveness arise from carefully orchestrated interactions between individual building blocks.^[3]^ Designing synthetic analogs with similar precision, however, remains challenging, as it requires fine-tuning of intermolecular forces that govern folding, assembly, and long-term stability.^[4]^ A variety of encapsulation matrices have been explored to stabilize sensitive molecules; however, among the most promising strategies to stabilize and spatially organize proteins is the use of metal–organic frameworks (MOFs), highly porous, modular architectures assembled from inorganic nodes and organic linkers.^[5]^ Their high surface area, tunable chemistry, and mild aqueous synthesis conditions enable biomacromolecule encapsulation with minimal structural disruption. Recent advances in MOF engineering have increasingly embraced biomimetic strategies not only for structural control but also for functional integration. For example, 1D MOF-based silk-like materials have been fabricated by harnessing directional MOF growth to form fibrous architectures with mechanical flexibility and hierarchical order, mimicking natural silk assembly.^[6a]^ Similarly, 2D MOF-based films have been developed by confining MOF crystallization at interfaces, enabling the formation of thin, flexible, and highly porous sheets suitable for membrane separation and sensing applications.^[6b]^ These examples highlight the versatility of biomolecular templates and interfaces in directing MOF morphology and functionality. Particularly, 3D zeolitic imidazolate frameworks (ZIFs) offer biocompatible synthesis pathways that support room-temperature self-assembly, preserving the functional conformation of sensitive proteins.^[5c, 7]^ Despite this promise, the mechanisms governing protein@MOF assembly at the molecular interface remain poorly understood.^[8]^ Key to this process is the interfacial environment, where electrostatic forces, hydrogen bonding, and hydrophobic interactions dictate nucleation, entrapment efficiency, and protein conformation.^[7a, 9]^ Yet, a major gap in the field remains in understanding how to rationally design the interfacial interactions between proteins and MOFs, and in elucidating the roles of surfactants and lipid-based amphiphiles in modulating protein–MOF interfaces.

In this study, we introduce a biomimetic strategy for engineering the protein@MOF architectures through surfactant-mediated interfacial design (Figure 1a and Figure S1). Inspired by the stabilizing role of lipid membranes in preserving protein structure and function, this approach leverages the amphiphilic and electrostatic properties of surfactants to modulate the interfacial environment of proteins during MOF formation. We anticipate that mimicking such membrane-like flexibility at the protein–MOF boundary can improve structural retention and functional performance of encapsulated proteins. By tailoring the nature of the surfactant at the protein–MOF interface, we aim to control nucleation behavior, improve protein entrapment, and maintain structural integrity of the protein within the crystalline matrix. To implement this new concept, we selected a diverse panel of surfactants, including non-ionic and ionic species, with varied hydrophobic tail structures and headgroup chemistries that allow interrogation of multiple interaction modes at the interface. This approach enables a systematic investigation into how surfactant properties influence protein@MOF assembly. Through combined experiments and molecular dynamics (MD) simulations, we demonstrate the feasibility of this strategy and uncover the main guiding principles for interfacial design. The key innovations in this work are twofold: (1) it significantly deepens our molecular-level mechanistic understanding of the assembly of robust protein@MOF composites, and (2) it develops a practical and general framework for designing new bioinorganic materials with superior protein stability, which can be applied in drug-delivery, biocatalysis, biosensing, and membrane protein stabilization.

**Figure 1.**
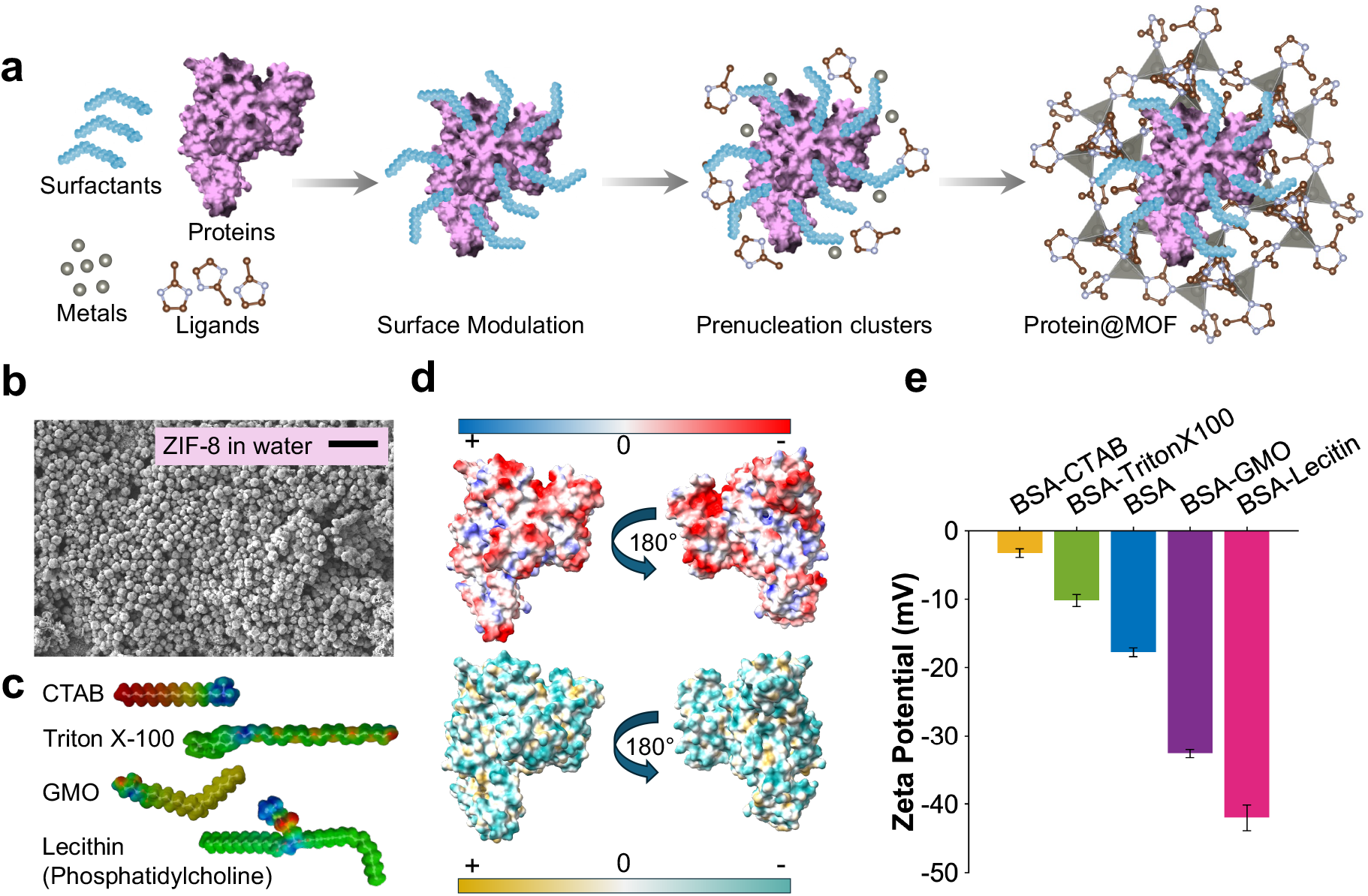
Synthesis and characterization of BSA@MOF. a) Schematic representation of MOF growth on the BSA surface in the presence of surfactants. b) Scanning electron microscopy (SEM) image of rhombic ZIF-8 formed in water, scale bar: 10 µm. c) The structure of four non-ionic and ionic surfactants; blue: highly hydrophilic (polar) regions, green: intermediate or neutral hydrophobicity, yellow/orange/red: increasingly hydrophobic (nonpolar) regions. d) Schematic representation of BSA’s electrostatic potential, red: negative potential, white: zero, blue: positive potential, and hydrophobicity distribution, dark cyan for most hydrophilic and dark goldenrod for most hydrophobic, PDB ID: 4F5S. e) Comparison of the zeta potentials for native BSA and BSA treated with various surfactants.

## 2. Results and Discussion

Within cellular environments, multivalency arising from tandem binding sites and repetitive motifs can drive the formation of biomolecular condensates,^[10]^ thereby contributing to the maintenance of intracellular crowding as well as structural and functional organization.^[11]^ Previous work by Patterson’s, Ge, and Ouyang’s groups revealed that in the coprecipitation process of protein@MOF, the metal ions and organic ligands could enhance multivalent interactions with proteins.^[12]^ Building on this initial understanding of protein@MOF formation, we sought to investigate how the surface characteristics of the protein contribute to the assembly process. In physiological systems, cell membranes predominantly adopt lamellar liquid crystalline phases, where phospholipids self-assemble into stable bilayers in aqueous environments.^[13]^ However, in specialized organelles such as microsomes and mitochondria, non-lamellar liquid crystalline structures characterized by higher curvature and dynamic flexibility are more prevalent.^[14]^ Inspired by the critical functional roles of membrane proteins, which emerge largely through their interactions with these diverse lipid architectures,^[15]^ we hypothesized that introducing surfactants at the protein@MOF interface could similarly modulate interfacial properties. This adaptive interfacial environment may better accommodate the structural requirements of proteins, enhancing their compatibility with the MOF scaffold and thereby preserving both their functionality and structural integrity. To verify this hypothesis, protein@MOF was synthesized via a one-pot coprecipitation process. Bovine serum albumin (BSA), a widely studied globular protein known for its stability and versatility, was adopted as the biomacromolecule model. Its well-characterized surface chemistry and amphiphilic nature make it an excellent model system for studying surfactant-mediated interfacial modulation. In a typical encapsulation process, BSA was initially dispersed in water, followed by the addition of the surfactant solution to alter the protein’s interfacial properties. Subsequently, zinc nitrate and 2-methylimidazole (HmIM) were introduced to initiate the biomimetic mineralization of ZIF-8 around BSA, leading to the formation of BSA@MOF (Figure 1a-b). In this process, the interfacial interactions between the protein and MOF precursors, including electrostatic forces, hydrophobic/hydrophilic interactions, and potential enzyme reorientation, play key roles in directing nucleation and crystallization pathways.^[16]^

To systematically investigate how surfactants modulate BSA surface properties, we tested different concentrations of the cationic surfactant, cetyltrimethylammonium bromide (CTAB), along with three non-ionic surfactants, including glycerol monooleate (GMO), lecithin, and Triton X-100 (Figure 1c). An initial surfactant concentration screen (70, 150, 300 µM) identified 70 µM as optimal for protein@MOF formation. These surfactants are hypothesized to modulate the surface charge and hydrophobicity index of BSA (Figure 1d), thereby impacting the nucleation and growth behavior of BSA@MOF composites, as well as their resulting structural stability.^[17]^

The zeta potential (ζ) of BSA was measured in the presence of each surfactant to quantify the effects on protein surface charge and potential interfacial restructuring (Figure 1e). Our findings revealed that the addition of surfactants significantly altered the zeta potential of BSA, indicating substantial modulation of interfacial charge interactions (Figure 1e). At pH 7, native BSA, with an isoelectric point (pI) of 4.9, exhibited a ζ potential of −17.8± 1.3 mV (Figure 1e), consistent with its anionic nature due to the deprotonation of carboxylic acid groups (glutamate and aspartate).^[18]^ The addition of non-ionic lipid-based surfactants, such as GMO and soy lecithin, led to a significant increase in negative charge density, lowering the ζ potential to −32.6± 1.2 mV and −42.0± 3.7 mV, respectively (Figure 1e). This suggests that GMO and lecithin contribute to BSA stabilization by increasing electrostatic repulsion at the protein interface, likely facilitated by hydrogen bonding and non-covalent interactions, which help maintain structural integrity and prevent aggregation.^[14, 19]^ GMO is a glycerol fatty acid ester characterized by a *cis* double bond at the C9 position, comprising a hydrophilic glycerol head group capable of hydrogen bonding in aqueous environments and a hydrophobic acyl tail, rendering it highly amphiphilic (Figure 1c).[^14, 20^] In our formulation process, GMO was initially dissolved in ethanol and subsequently incorporated into the aqueous BSA solution. Given that GMO readily forms a cubic liquid crystalline phase (specifically, the diamond D phase) in environments containing greater than 40% water,^[14]^ it is plausible that such non-lamellar structural organization at the protein interface contributed to the observed stabilization due to the local restructuring of the BSA hydration layer. The presence of these non-lamellar phases may provide a dynamic, flexible interfacial environment that better accommodates the BSA molecules.

Similarly, lecithin, a natural mixture of polar lipids (glycolipids, phospholipids) and neutral lipids (triglycerides), may have facilitated a local restructuring of the BSA hydration layer due to its key role in the self-assembly of lyotropic liquid crystalline phases,^[21]^ contributing to the pronounced decrease in zeta potential (Figure 1e). In contrast, Triton X-100, another non-ionic surfactant, and CTAB, a cationic surfactant, did not promote non-lamellar phase transitions, but instead reduced the magnitude of BSA’s surface charge. Triton X-100 caused a moderate increase in zeta potential to −10.2± 1.7 mV, while CTAB induced the most significant shift, raising it to −3.3± 1.3 mV (Figure 1e). As a bulky ethoxylated surfactant, Triton X-100 likely intercalated its hydrophobic tail into exposed non-polar regions of BSA, disrupting electrostatic interactions without forming an organized interfacial phase.^[22]^ Meanwhile, CTAB effectively neutralized BSA’s negative surface charge through strong electrostatic attraction between its positively charged ammonium groups and BSA’s anionic residues, leading to a significant charge compensation effect.^[23]^

To further investigate the structural stability of BSA in the presence of surfactants, we employed far-UV circular dichroism (CD) spectroscopy, which is sensitive to the conformational state of the protein backbone.^[24]^ CD spectra revealed that BSA retains a predominantly α-helical structure under all tested conditions, as evidenced by the characteristic negative ellipticity bands at 208 and 222 nm and a positive band near 193 nm (Figure 2a), consistent with its native conformation.^[25]^ Quantitative secondary structure analysis using the BeStSel algorithm indicated that native BSA exhibits approximately 70% α-helical content, in agreement with its previously reported crystal structure, PDB ID: 4F5S (Figure S2).^[26]^ Upon addition of surfactants, distinct trends emerged in BSA’s secondary structure content. GMO led to the most pronounced increase in α-helicity, raising it to 82%, followed by lecithin and Triton X-100, both of which enhanced helical content to 75% (Figure 2b). This trend closely mirrors the zeta potential results, where GMO and lecithin significantly increased BSA’s negative surface charge. The enhancement in α-helical structure in the presence of GMO is likely attributable to its ability to form non-lamellar cubic phases, which provide a dynamic, hydrated, and hydrogen-bond-rich interfacial environment that supports native protein folding.^[14]^ Lecithin, which also promotes lyotropic liquid crystalline phase formation,^[21]^ appears to exert a similar but slightly less pronounced stabilizing effect. Triton X-100, although non-ionic, did not support organized interfacial architectures, and its moderate increase in α-helicity may result from partial shielding of hydrophobic regions without strong structural reinforcement.^[22]^ In contrast, CTAB, a cationic surfactant, decreased α-helical content to 63%, suggesting partial unfolding or conformational destabilization (Figure 2b). This loss of structure correlates with CTAB’s significant neutralization of BSA’s surface charge and highlights the disruptive effect of strong electrostatic binding on protein folding.^[23]^

**Figure 2.**
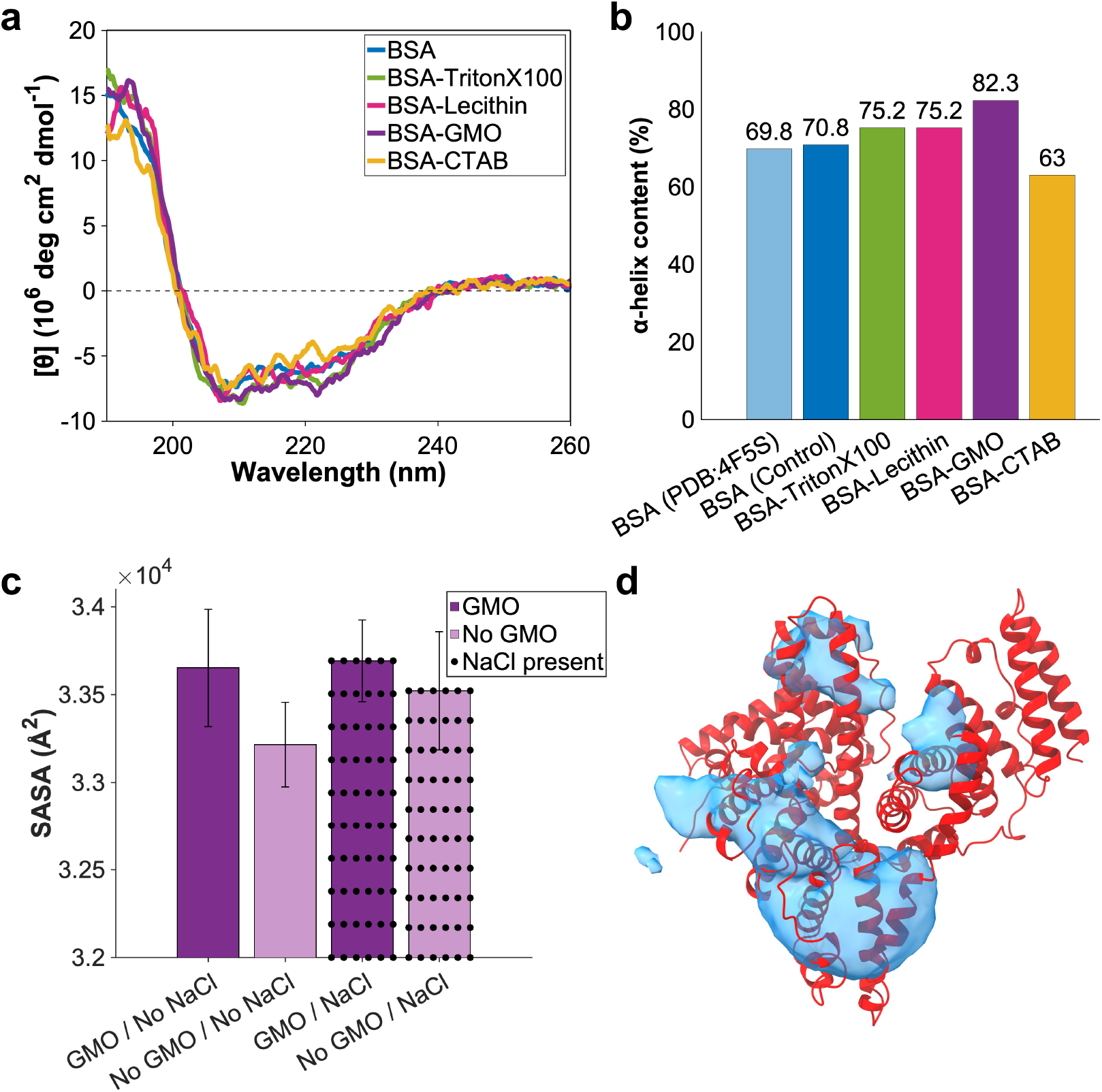
a) Circular dichroism (CD) spectra of native BSA compared to BSA treated with various surfactants. b) Percentage of α-helix content in native BSA and BSA treated with surfactants. (c) Solvent-accessible surface area (SASA) values for BSA in the presence and absence of GMO averaged over six independent MD simulations with and without NaCl concentration. d) Spatial distribution of the surfactant head groups around BSA, represented as an isosurface with averaged occupancy of 0.005. The protein is displayed in red, and the blue isosurface highlights regions around the protein visited most frequently by the head groups of GMO molecules. The occupancy isosurface was generated by averaging the center-of-mass positions of the GMOs’ head groups across six independent MD trajectories without NaCl concentration.

Motivated by these findings, we sought to examine the molecular interactions at the BSA– surfactant interface using all-atom molecular dynamics (MD) simulations in the presence and absence of the GMO molecules and a 0.15 M NaCl concentration, with particular focus on GMO’s stabilizing effect on protein structure. These simulations enabled us to identify the protein residues that interact most strongly with surfactant molecules and to evaluate how these interactions are modulated by ionic strength. Moreover, solvent-accessible surface area (SASA) values and occupancy maps were calculated (Figure 2, c-d). To identify preferential binding sites, we calculated the spatial distribution of GMO headgroups around the protein, which is represented as isosurfaces with occupancy beyond 0.005 (Figure 2d). Additionally, we calculated the conformational factor (*P*_*i*_, with *i* being the residue number) for each residue, defined as the relative contact frequency between GMO headgroups and individual amino acids across all trajectories. Residues with *P*_*i*_ ≫ 1 were considered to exhibit above-average interaction with GMO.

Both hydrophobic and hydrophilic residues showed significant interactions with the surfactant (Figure S3 and S4), indicating that GMO binding is not limited to a single residue type. We identified residues with *P*_*i*_ ≫ 1 that consistently appeared across simulation setups under both 0 M and 0.15 M NaCl conditions, including Glu186, Lys187, Phe205, Phe227, Thr231, Asp323, Ala324, Lys350, Arg435, Lys439, and Tyr451 (Table S1). These residues encompass a diverse set of physicochemical classes, acidic (Glu, Asp), basic (Lys, Arg), polar (Thr, Tyr), aromatic (Phe), and aliphatic (Ala). They are predominantly hydrophilic residues, highlighting GMO’s ability to engage in both electrostatic and hydrophobic interactions. This molecular versatility is consistent with earlier experimental data that demonstrated GMO enhanced the magnitude of BSA’s negative zeta potential (Figure 1e) and increased α-helicity (Figure 2b), potentially through hydrogen bonding with polar residues and electrostatic interactions with charged side chains. Moreover, interactions with hydrophobic and aromatic residues (e.g., Phe227, Tyr451) suggest that GMO’s acyl tail can embed into local nonpolar pockets on the protein surface, stabilizing its conformation (Table S1 and Figure 2, c-d). The presence of 0.15 M NaCl reduced the *P*_*i*_ values of several high-affinity residues, indicating that ionic strength could modulate GMO–BSA interactions by screening electrostatic forces (Table S1). This trend was also reflected in a reduced change in SASA in the salt-containing system (Figure 2c), further supporting the role of ionic strength in reducing surfactant–protein association.

The residues with highest *P*_*i*_ values clustered within domains IIA and IIIA of BSA (Figure 3, a-b), regions known to contain flexible loops and hydrophobic cavities that facilitate ligand binding.^[27]^ This observation suggests that GMO does not bind randomly but instead localizes near functionally relevant surface regions. These regions are characterized by favorable local polarity, side chain flexibility, and surface accessibility, which together promote stable interactions. The spatial distribution of GMO headgroups supports this interpretation and clarifies how GMOs and similar surfactants can influence protein stability and solubility through targeted interactions with specific surface regions. The occupancy isosurface reveals regions where GMO headgroups have high occupancy. These regions were around domains IIA and IIIA (Figure 2d) and closely correspond to the residue-level interaction patterns identified by the *P*_*i*_ analysis. To assess whether GMO binding altered BSA’s solvent exposure, we compared the SASA of BSA with and without surfactants under both ionic conditions (Figure 2c). The presence of GMO increased SASA relative to the control, indicating partial remodeling of the protein surface and modest conformational rearrangements that potentially perturb protein–surfactant interactions and interfacial hydration (Figure 2c). In contrast, the inclusion of 0.15 M NaCl attenuated this effect, leading to a smaller SASA increase and a reduction in overall GMO–protein interaction (Figure 2c).

**Figure 3.**
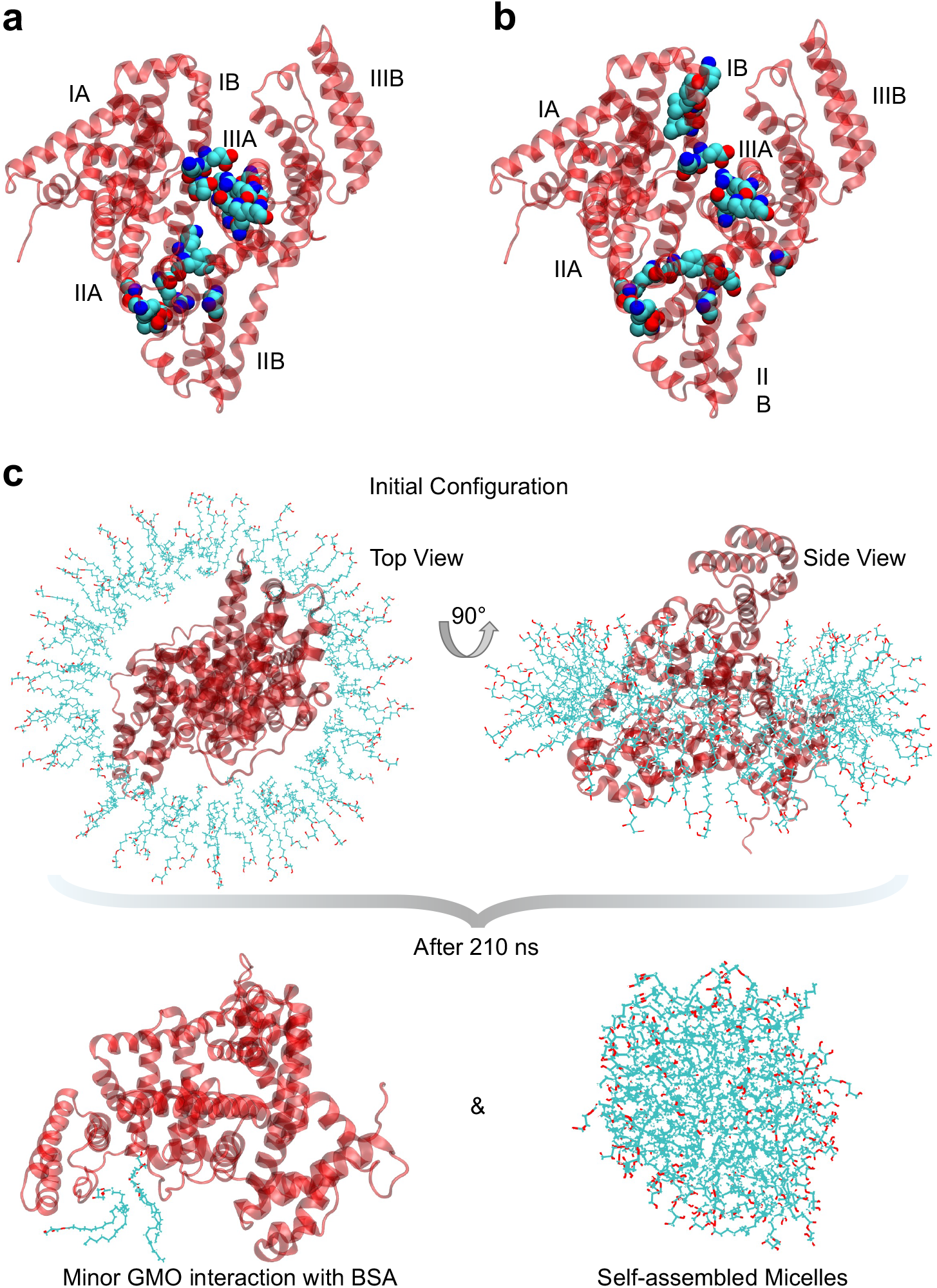
a) BSA residues with *P*_*i*_ < 1 with 0 M NaCl concentration. b) BSA residues with *P*_*i*_ < 1 in the presence of 0.15 M NaCl. The residues Glu186, Lys187, Phe205, Phe227, Tyr231, Asp323, Ala324, Lys350, Arg435, Lys439, and Tyr451 consistently exhibited above-average interactions with GMO across all simulation setups, both with and without NaCl. These residues represent the top 20 *P*_*i*_ values, averaged across all six independent simulations. All *P*_*i*_ values exceed 5. The BSA domains are labeled. c) Initial configuration of BSA and 100 GMO molecules at 0 ns, shown from top and side views, and final structure after 210 ns of MD simulation. At this higher concentration, GMO molecules exhibit reduced interaction with BSA.

Consistent with our above-mentioned analysis, this indicates that the salt concentration also plays an important role in modulating the interaction between GMO and the BSA. Moreover, the MD simulation results indicate that at higher GMO concentrations, GMO interacts less effectively with BSA (Figure 3c). In the system containing 100 GMO molecules, the surfactants aggregate to form a complete micelle, resulting in predominant GMO-GMO interactions rather than interactions with BSA (Figure 3c). This self-assembly behavior is potentially driven by the inherent amphiphilic property of GMO molecules which clusters their hydrophobic tails in the inner core of the micelle and exposes their headgroups to the aqueous environment. Consequently, fewer free GMO molecules are available to interact with the protein surface.

These findings are particularly important for the design of protein@MOF composites, where interfacial stabilization and controlled nucleation are critical for structural retention and encapsulation efficiency. Ge and co-workers discovered that during the biomineralization process of protein@MOF, protein molecules form clusters with MOF precursors, contributing to structural and functional integrity and overall activity retention by sacrificing protein molecules at the cluster surface.^[12c]^ We speculate that lipid-based non-ionic surfactants such as GMO may reduce the extent of chemical destruction by local MOF precursors, forming a protective layer at the protein-MOF interface while wrapping proteins inside. Previous studies have also highlighted the role of negatively charged carboxylate groups in amino acid residues in facilitating MOF nucleation and accelerating framework growth around encapsulated biomolecules.^[5c, 7a, 28]^ Additionally, hydrophilicity and hydrophobicity at the protein–MOF interface have been shown to significantly influence enzyme activity and stability upon encapsulation. For instance, hydrophilic MOFs such as ZIF-90 and MAF-7 effectively preserve enzymatic function by protecting proteins from denaturing conditions, whereas encapsulation in hydrophobic MOFs like ZIF-8 often results in enzyme inactivation or minimal activity.^[9a]^ This suggests that hydrophilic enzymes favor hydrophilic MOFs, leading to better stabilization and retention of catalytic activity. However, contradictory findings indicate that hydrophobic interactions can also enhance enzyme stability. For example, MD simulations of cutinase-encapsulated MOF-74 (IRMOF-74-VI) demonstrated that hydrophobic amino acid residues (e.g., Arg) formed hydrogen bonds and salt bridges with hydrophobic linkers, maintaining structural integrity even at elevated temperatures (500 K).^[9b]^ To evaluate the functional impact of surfactants on biomineralization, we next examined how surfactants at the interface of proteins modulate the growth kinetics of ZIF-8 on the BSA surface (Figure 4a, S5, S6, and Table S2). Using time-resolved dynamic light scattering (DLS), we monitored the evolution of particle size and fitted the resulting curves to an exponential model to extract initial growth rates (Figure S5, S6, and Table S2). Among the surfactants tested, lecithin resulted in the most pronounced increase in MOF growth rate, reaching 35.50 ± 4.84 nm/s. GMO followed closely with a rate of 34.30 ± 5.99 nm/s. Triton X-100 produced a moderate growth rate of 23.28 ± 8.31 nm/s, which was not significantly different from the BSA control (27.05 ± 4.61 nm/s). Notably, CTAB markedly inhibited MOF formation, reducing the initial growth rate to 10.20 ± 3.20 nm/s (Figure 4a, S5, S6, and Table S2). These results are consistent with the trend observed in zeta potential measurements (Figure 1e), where lecithin and GMO induced the largest increases in negative surface charge, potentially enhancing electrostatic interactions with zinc ions during ZIF-8 nucleation. In contrast, CTAB significantly neutralized the negative charge on BSA, thereby suppressing nucleation and growth, consistent with its destabilizing effect on BSA secondary structure (Figure 2a). In parallel, we assessed the efficiency of BSA encapsulation within ZIF-8 using a Bradford protein assay (Figure 4, b-c and Figure S7). Surprisingly, the BSA control without any surfactant exhibited the lowest encapsulation efficiency across all tested conditions. In contrast, the addition of lecithin or GMO increased protein loading by approximately 15–20%, while Triton X-100 and CTAB also produced moderate improvements (Figure 4c).

**Figure 4.**
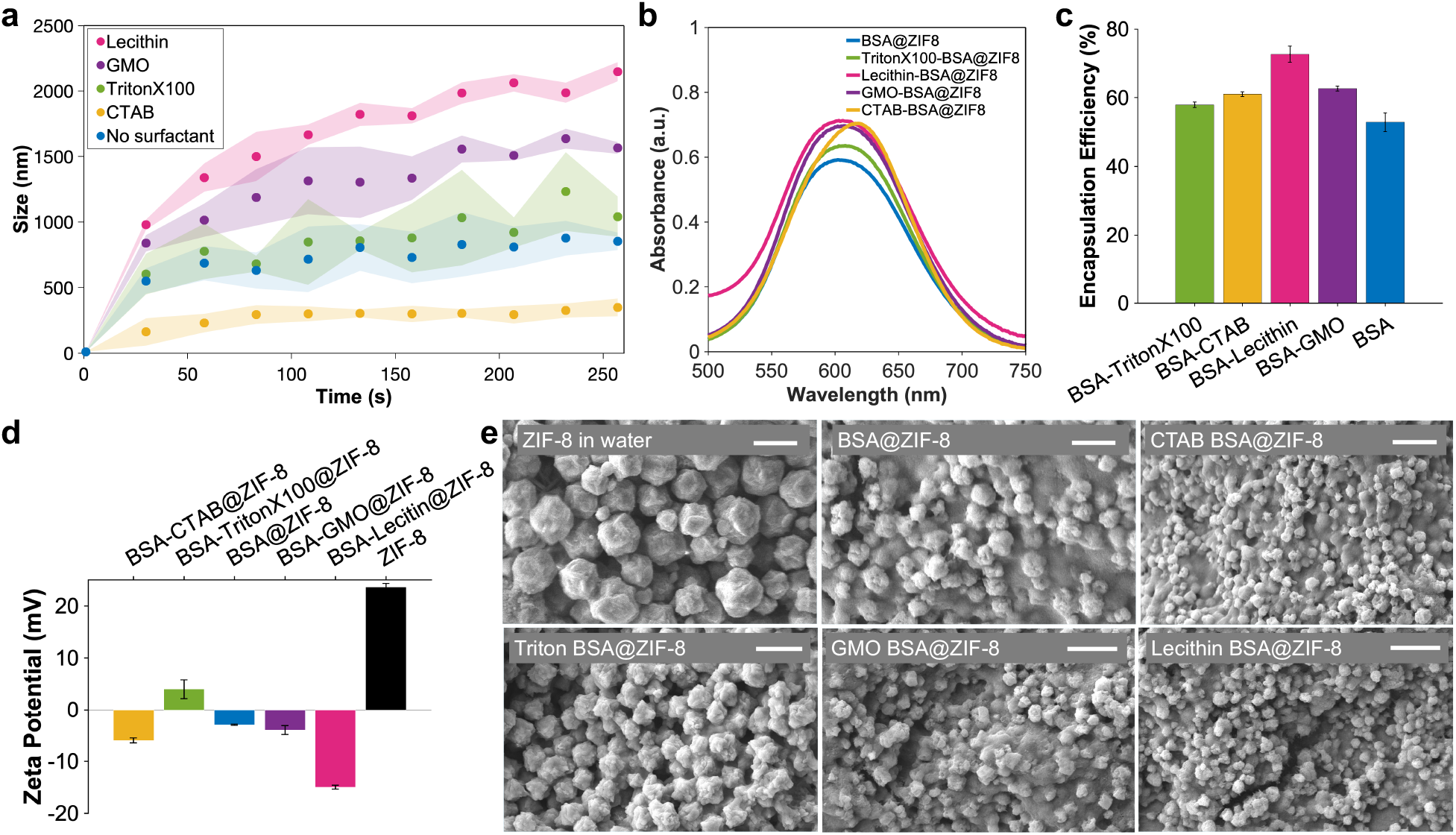
a) Scatter plots showing the size evolution of BSA@MOF composites over 250 seconds, with and without surfactants. b) Bradford assay measuring protein content in protein@MOF composites formed in the presence of different surfactants (*λ*_max_= 595 nm). c) Encapsulation efficiency (%) of BSA within MOFs synthesized with various surfactants. d) Zeta potential of BSA@MOF composites following synthesis. e) SEM images of rhombic BSA@MOF crystals formed in the presence of different surfactants. Scale bars: 2 µm.

The reduced encapsulation in the control condition may reflect limited interaction between native BSA and MOF precursors due to suboptimal surface presentation or repulsive electrostatic effects.^[9]^ Notably, the CD spectra of BSA in the presence of MOF precursors prior to ZIF-8 formation showed no significant conformational disruption relative to the BSA control (Figure S8), suggesting that changes in encapsulation efficiency are not due to unfolding but rather to interfacial compatibility. Additionally, elemental analysis revealed a clear increase in both carbon and sulfur content in the BSA@MOF samples compared to control MOF, further supporting successful encapsulation of the protein. The presence of sulfur, absent in pure ZIF-8, is attributed to sulfur-containing amino acids such as cysteine and methionine, providing direct evidence of protein incorporation within the MOF framework (Table S3 and Figure S9).^[12c]^ These findings further support the notion that amphiphilic surfactants, particularly lecithin and GMO, not only enhance MOF nucleation but also improve protein entrapment through modulation of interfacial structure and electrostatics.

To investigate the impact of surfactants on the surface properties of the resulting BSA@MOF, we measured the zeta potential of the particles after MOF formation (Figure 4d). Control ZIF-8 exhibited a highly positive zeta potential (+23.6 ± 1.5 mV), as expected due to its zinc-rich framework. Interestingly, the presence of BSA and surfactants during the encapsulation process decreased the surface charge. Lecithin produced BSA@MOF particles with the most negative zeta potential of −15.0 ± 0.7 mV, suggesting strong interfacial interactions and effective surface modification (Figure 4d). The zeta potential results correlate well with particle morphology observed via scanning electron microscopy (SEM). Control ZIF-8 displayed the largest size, averaging approximately 2 µm (Figure 4e). In contrast, the addition of surfactants during synthesis led to a marked reduction in particle dimensions. Lecithin yielded the smallest particle size (400 nm), followed by GMO (500 nm), CTAB (600 nm), and Triton X-100 (1 µm) (Figure 4e). SEM images showed similar rhombic dodecahedral structures with a rough surface. These results suggest that surfactants, particularly lecithin and GMO, significantly influence nucleation and growth processes, promoting the formation of smaller, more compact MOF structures.

In addition, 77K N_2_ sorption test was conducted for ZIF-8 and protein@ZIF-8 in the presence and absence of surfactants to evaluate the textural properties of composites and the effect of protein encapsulation and surfactants on pore structure (Figure S10, a-b). The data obtained from multipoint Brunauer–Emmett–Teller (BET) and Density Functional Theory (DFT) analysis are summarized in Table S4. Interestingly, the pore width of ZIF-8 remains constant at approximately 17.2 Å across all samples (Figure S10 and Table S4), which is within the reported literature range and confirms that protein encapsulation does not disrupt the intrinsic pore size or geometry of the framework. The high pore width would potentially reduce the mass transfer resistance of small molecules for biocatalysis applications. The surface area of the pristine ZIF-8 measured ∼2060 m^2^/g, which confirms the high crystallinity and efficient activation during sample preparation. Upon protein encapsulation, a consistent decrease in surface area is observed across all samples with and without surfactant, indicating successful incorporation of the protein within the porous matrix (Table S4). Interestingly, variations among the surfactant-assisted composites reveal that different surfactants influence the surface area to varying degrees. For example, BSA-Lecithin@ZIF-8 exhibit lower surface areas than BSA@ZIF-8, which could be due to the stronger dual interactions of lecithin with both protein and the ZIF-8 interface. This event can potentially enhance protein encapsulation in MOF using lecithin, which is in agreement with the encapsulation efficiency data presented in Figure 4C, and lead to greater pore filling and lower surface areas (Table S4).

Powder X-ray diffraction (XRD) patterns confirmed the crystalline structure of ZIF-8 in the BSA@ZIF-8, both before and after protein encapsulation and in the presence of surfactants (Figure 5a). All BSA@ZIF-8 samples, with or without surfactants, exhibited diffraction patterns consistent with the characteristic ZIF-8 framework (Figure 5a). These findings are consistent with SEM, which showed a similar rhombic dodecahedral morphology for both ZIF-8 and BSA@ZIF-8 (Figure 4e). Thermogravimetric analysis (TGA) curves showed distinct weight loss around 300 °C for free BSA and BSA@ZIF-8, indicating the successful incorporation of BSA into the MOF. The increased char yield from 15% for free BSA to 38% for BSA-GMO@ZIF-8 suggests enhanced thermal stability of BSA upon encapsulation (Figure 5b and Table S5). Previous studies have demonstrated that encapsulating biomacromolecules within MOFs can significantly enhance their stability under harsh conditions and extend their storage time.^[28]^ For example, biomimetic mineralization using MOFs has been shown to protect proteins, enzymes, and DNA from denaturation by forming a crystalline exoskeleton under mild, physiological conditions. These protective shells preserve bioactivity even after exposure to extreme thermal and chemical stressors, including boiling in organic solvents such as dimethylformamide and heating up to 80 °C, which would normally inactivate free proteins.^[28]^ These findings support the idea that MOF-based encapsulation can enhance protein stability and function, aligning with our design strategy for surfactant-mediated protein@MOF assemblies. Attenuated total reflection Fourier transformed infrared spectroscopy (ATR-FTIR) confirmed the encapsulation of BSA by the appearance of the amide band in 1700−1500 cm^−1^ region (Figure 5c).

**Figure 5.**
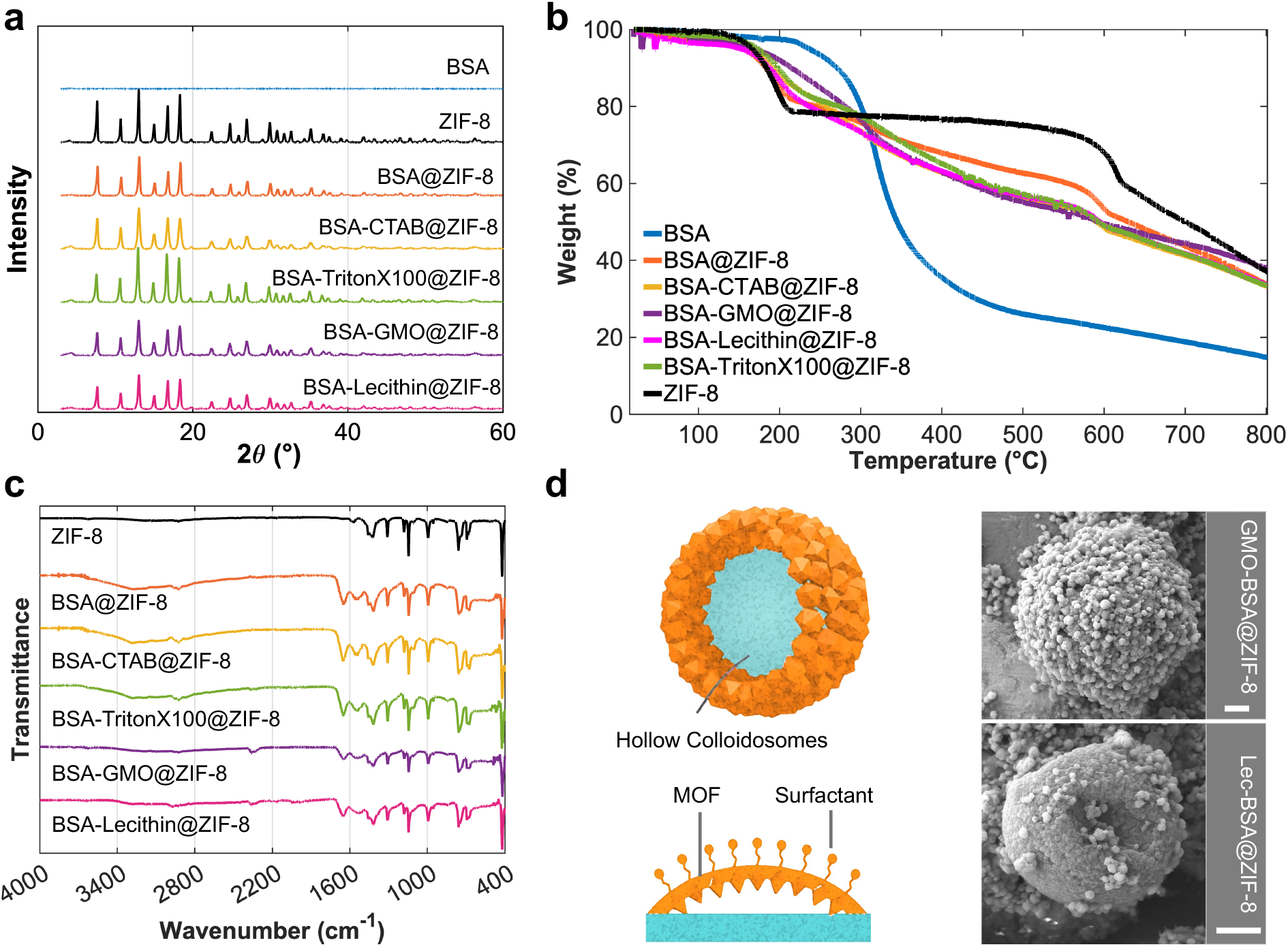
a) PXRD patterns of BSA@MOF composites showing consistent crystallinity in the presence of various surfactants. b) TGA curves of free BSA and BSA@MOF composites synthesized with different surfactants. c) ATR-FTIR spectra of protein@MOF composites formed in the presence of various surfactants. d) Formation of hollow colloidosomes assembled from individual rhombic BSA@MOF particles in the presence of GMO and lecithin, scale bar: 2 µm.

Additionally, we observed the self-assembly of colloidosomes approximately 10 µm in size in both BSA-GMO@ZIF-8 and BSA-lecithin@ZIF-8 samples (Figure 5d). This indicates the ability of GMO and lecithin to facilitate the formation of Pickering emulsion droplets,^[29]^ with BSA@ZIF-8 particles decorating the colloidosome surface. Given that colloidosomes offer efficient substrate transport channels due to their core–shell architecture and can spatially organize catalytically active sites such as enzymes in defined orientations,^[29]^ these findings provide valuable insight into a facile synthetic strategy for constructing multienzyme biocatalysts.

To validate the functional potential of our protein@MOF system for biocatalytic applications, we evaluated the enzymatic activity of horseradish peroxidase (HRP) encapsulated in ZIF-8 under different surfactant conditions (Figure S11, a-b). In this assay, hydrogen peroxide was provided as the oxidant and o-phenylenediamine (OPD) was used as the substrate. HRP catalyzes the oxidation of OPD by hydrogen peroxide, a prototypical peroxidase reaction that involves O–O bond activation followed by oxidation of aromatic C–H bonds on OPD, producing a yellow oxidized product with strong absorbance at 450 nm. Fresh free HRP was used as a baseline with 100% activity. Direct encapsulation into ZIF-8 without surfactants yielded only ∼13% of native activity, consistent with prior reports due to potential restricted substrate accessibility. In contrast, surfactant-assisted encapsulation markedly improved activity retention: HRP–lecithin@ZIF-8 preserved ∼62% of native activity, while HRP–GMO@ZIF-8 retained ∼16% (Figure S11b). These results highlight the beneficial role of surfactants in preserving enzyme functionality and support the promise of this strategy for application in biocatalysis.

Moreover, protein-GMO@MOF and protein-lecithin@MOF composites could be potentially used to stabilize the membrane proteins by imitating their dynamic interactions with lipids.^[15a]^ Thus, they show strong potential as versatile alternatives to lipid-based nanodiscs for stabilizing membrane proteins. This is especially important given the ongoing challenge of preserving membrane protein integrity in cell-free environments, where native lipid bilayers are absent, and denaturation is common.^[30]^ Unlike traditional nanodiscs, which often exhibit limited long-term stability and incompatibility with solid-state or heterogeneous environments,^[31]^ protein@MOF composites with integration of lipid-based non-ionic surfactants may offer a more robust and adaptable platform. This has significant implications for pharmaceutical research on sensitive proteins,^[32]^ where approximately 60% of drug targets are membrane-associated proteins.^[15b, 15c, 33]^ Developing stable, bioinspired matrices that preserve protein conformation and activity could open new pathways for drug screening and delivery, biocatalysis and biosensing.

## 3. Conclusion

In summary, this study establishes a mechanistic framework for understanding how surfactant-mediated interfacial design governs the encapsulation of proteins within metal–organic frameworks. By employing a combination of biochemical assays, spectroscopy, microscopy, and MD simulations, we reveal how surfactants modulate protein–MOF interactions through alterations in electrostatic potential, hydrophobicity, and protein surface presentation. Our findings show that lipid-based non-ionic surfactants, particularly GMO and lecithin, promote favorable interfacial environments that preserve protein secondary structure, enhance encapsulation efficiency, and enable the formation of colloidosome-like architectures. Importantly, domain-specific binding patterns observed in simulations support a model where localized, stabilizing interactions between surfactant molecules and protein residues guide MOF nucleation and particle organization. These insights contribute to a clearer understanding of how proteins behave at the interface and underscore the role of molecular design in optimizing protein@MOF formation. Beyond advancing the fundamental mechanistic understanding of the assembly of biomolecular inorganic materials, this work offers a versatile and practical strategy for engineering such materials with improved retention of structure and function compared to traditional techniques based on nanodiscs. Thus, it opens new doors to future applications in drug delivery, biocatalysis, biosensing, and membrane protein stabilization.

## Acknowledgements

The authors thank Bo Zhao, Reagan Hudson, Jeshua Podliska, Anthony Cozzolino, Hariharan Parameswaran, and Bailey Bouley for their assistance with data acquisition for SEM, PXRD, FTIR, TGA, CD, BET, and elemental analysis. We acknowledge support from the College of Arts & Sciences Microscopy, X-ray Diffraction Facility, the computing facilities provided by the High-Performance Computing Center at TTU, and the Membrane Protein Laboratory Core at TTUHSC. We are grateful for the funding support from the ACS Petroleum Research Fund Award 69034-DNI3. R.L. and M.K. acknowledge funding support from the National Institutes of Health (R35GM150780). Molecular graphics and analyses performed with UCSF ChimeraX, developed by the Resource for Biocomputing, Visualization, and Informatics at the University of California, San Francisco, with support from National Institutes of Health R01-GM129325 and the Office of Cyber Infrastructure and Computational Biology, National Institute of Allergy and Infectious Diseases.

## Data Availability Statement

The supplementary information, raw data and custom analysis scripts are available upon request from the corresponding author.

## Table of Content

**Mechanistic Understanding of Protein–MOF Integration through Surfactant-Driven Interfacial Design**

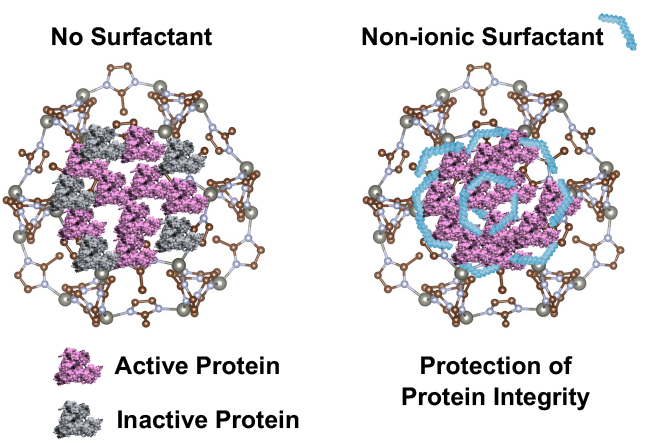

This study reveals how surfactant-driven interfacial design governs the assembly and stability of protein@MOF composites. Using lipid-based non-ionic surfactants, we modulate protein– MOF interactions to improve encapsulation efficiency and MOF crystallization. Molecular dynamics simulations uncover residue-specific interactions at the interface, providing mechanistic insight into the role of surfactants in guiding protein integration into porous frameworks.

## References

[1] V. Nguyen, C. Wilson, M. Hoemberger, J. B. Stiller, R. V. Agafonov, S. Kutter, J. English, D. L. Theobald, D. Kern, Science 2017, 355, 289.

[2] a)J. Wang, Y. Li, G. Nie, Nature Reviews Materials 2021, 6, 766; b)Y. Lu, A. A. Aimetti, R. Langer, Z. Gu, Nature Reviews Materials 2016, 2, 1; c)J. Han, M. Fussenegger, Nature Reviews Materials 2025, 1; d)M. Wittwer, U. Markel, J. Schiffels, J. Okuda, D. F. Sauer, U. Schwaneberg, Nature Catalysis 2021, 4, 814.

[3] a)D. A. Fletcher, R. D. Mullins, Nature 2010, 463, 485; b)S. Keten, Z. Xu, B. Ihle, M. J. Buehler, Nature materials 2010, 9, 359; c)J. C. van Hest, D. A. Tirrell, Chemical communications 2001, 1897; d)A. M. Kushner, Z. Guan, Angewandte Chemie International Edition 2011, 50, 9026; e)M. J. Buehler, Y. C. Yung, Nature materials 2009, 8, 175.

[4] F. Barthelat, Z. Yin, M. J. Buehler, Nature Reviews Materials 2016, 1, 1.

[5] a)M. Eddaoudi, D. B. Moler, H. Li, B. Chen, T. M. Reineke, M. O’keeffe, O. M. Yaghi, Accounts of chemical research 2001, 34, 319; b)O. M. Yaghi, M. O’Keeffe, N. W. Ockwig, H.K. Chae, M. Eddaoudi, J. Kim, Nature 2003, 423, 705; c)H. Wang, L. Han, D. Zheng, M. Yang, Y. H. Andaloussi, P. Cheng, Z. Zhang, S. Ma, M. J. Zaworotko, Y. Feng, Angewandte Chemie International Edition 2020, 59, 6263; d)Y. Feng, H. Wang, S. Zhang, Y. Zhao, J. Gao, Y. Zheng, P. Zhao, Z. Zhang, M. J. Zaworotko, P. Cheng, Advanced Materials 2019, 31, 1805148; e)R. Ravanfar, T. A. Comunian, A. Abbaspourrad, Food Hydrocolloids 2018, 81, 419–428; f)R. Ravanfar, T. A. Comunian, R. Dando, A. Abbaspourrad, Food chemistry 2018, 241, 460–467. g) R. Ravanfar, G. B. Celli, A. Abbaspourrad, ACS Applied Materials & Interfaces 2018, 10 (6), 6046–6053, h) R. Ravanfar, C. J. Bayles, A. Abbaspourrad, Crystal Growth & Design 2020, 20 (3), 1673–1680, i) R. Ravanfar, A. Abbaspourrad, ACS Applied Materials & Interfaces 2019, 11 (42), 39376–39384.

[6] a)M. Cheng, M. Cao, X. H. Bu, Advanced Engineering Materials 2025, 2500592; b)M. Cheng, H. Bai, X. Wang, Z. Chang, M. Cao, X. H. Bu, Advanced Materials 2024, 36 (48), 2411680.

[7] a)G. Chen, S. Huang, X. Kou, S. Wei, S. Huang, S. Jiang, J. Shen, F. Zhu, G. Ouyang, Angewandte Chemie International Edition 2019, 58, 1463; b)J. B. Bailey, F. A. Tezcan, Journal of the American Chemical Society 2020, 142, 17265; c)G. Chen, S. Huang, X. Kou, F. Zhu, G. Ouyang, Angewandte Chemie 2020, 132, 14051; d)S. Huang, X. Kou, J. Shen, G. Chen, G. Ouyang, Angewandte Chemie International Edition 2020, 59, 8786.

[8] a)W. Liang, P. Wied, F. Carraro, C. J. Sumby, B. Nidetzky, C.-K. Tsung, P. Falcaro, C. J. Doonan, Chemical reviews 2021, 121, 1077; b)W. Liang, R. Ricco, N. K. Maddigan, R. P. Dickinson, H. Xu, Q. Li, C. J. Sumby, S. G. Bell, P. Falcaro, C. J. Doonan, Chemistry of Materials 2018, 30, 1069; c)R. Dhaoui, S. L. Cazarez, L. Xing, E. Baghdadi, J. T. Mulvey, N. S. Idris, P. J. Hurst, M. P. Vena, G. D. Palma, J. P. Patterson, Advanced Functional Materials 2024, 34, 2312972.

[9] a)W. Liang, H. Xu, F. Carraro, N. K. Maddigan, Q. Li, S. G. Bell, D. M. Huang, A. Tarzia, M. B. Solomon, H. Amenitsch, Journal of the American Chemical Society 2019, 141, 2348; b)T. Tuan Kob, M. Ismail, M. Abdul Rahman, K. E. Cordova, M. Mohammad Latif, The Journal of Physical Chemistry B 2020, 124, 3678.

[10] S. Mehta, J. Zhang, Nature Reviews Cancer 2022, 22, 239.

[11] S. F. Banani, H. O. Lee, A. A. Hyman, M. K. Rosen, Nature reviews Molecular cell biology 2017, 18, 285.

[12] a)B. P. Carpenter, A. R. Talosig, J. T. Mulvey, J. G. Merham, J. Esquivel, B. Rose, A. F. Ogata, D. A. Fishman, J. P. Patterson, Chemistry of Materials 2022, 34, 8336; b)A. F. Ogata, A. M. Rakowski, B. P. Carpenter, D. A. Fishman, J. G. Merham, P. J. Hurst, J. P. Patterson, Journal of the American Chemical Society 2020, 142, 1433; c)Y. Liu, S. Cui, W. Ma, Y. Wu, R. Xin, Y. Bai, Z. Chen, J. Xu, J. Ge, Journal of the American Chemical Society 2024, 146, 12565; d)L. Tong, S. Huang, Y. Shen, S. Liu, X. Ma, F. Zhu, G. Chen, G. Ouyang, Nature Communications 2022, 13, 951.

[13] P. J. Collings, J. W. Goodby, Introduction to liquid crystals: chemistry and physics, Crc Press, 2019.

[14] S. Milak, A. Zimmer, International journal of pharmaceutics 2015, 478, 569.

[15] a)A.-E. Saliba, I. Vonkova, A.-C. Gavin, Nature Reviews Molecular Cell Biology 2015, 16, 753; b)J. P. Overington, B. Al-Lazikani, A. L. Hopkins, Nature reviews Drug discovery 2006, 5, 993; c)M. P. Wymann, R. Schneiter, Nature reviews Molecular cell biology 2008, 9, 162.

[16] a)B. P. Carpenter, A. R. Talosig, B. Rose, G. Di Palma, J. P. Patterson, Chem Soc Rev 2023, 52, 6918; b)B. P. Carpenter, A. R. Talosig, J. T. Mulvey, J. G. Merham, J. Esquivel, B. Rose, A. F. Ogata, D. A. Fishman, J. P. Patterson, Chem Mater 2022, 34, 8336; c)N. K. Maddigan, A. Tarzia, D. M. Huang, C. J. Sumby, S. G. Bell, P. Falcaro, C. J. Doonan, Chem Sci 2018, 9, 4217.

[17] a)J. Van Houten, R. C. Barberi, J. King, A. F. Ogata, Materials Advances 2024, 5, 5945; b)X. Xu, Y. Liu, Z. Guo, X. Z. Song, X. Qi, Z. Dai, Z. Tan, IET Nanobiotechnol 2020, 14, 595.

[18] K. Eisele, R. A. Gropeanu, C. M. Zehendner, A. Rouhanipour, A. Ramanathan, G. Mihov, K. Koynov, C. R. Kuhlmann, S. G. Vasudevan, H. J. Luhmann, Biomaterials 2010, 31, 8789.

[19] B. Xia, Y. Shen, R. Zhao, J. Deng, C. Wang, Food Hydrocolloids 2024, 155, 110168.

[20] C. V. Kulkarni, W. Wachter, G. Iglesias-Salto, S. Engelskirchen, S. Ahualli, Physical Chemistry Chemical Physics 2011, 13, 3004.

[21] E. Kozakiewicz^1^, D. Cossuta, Handbook of Molecular Gastronomy: Scientific Foundations, Educational Practices, and Culinary Applications 2021, 249.

[22] S. K. Singh, N. Kishore, The Journal of Physical Chemistry B 2006, 110, 9728.

[23] a)C. Zhou, H. Wang, H. Bai, P. Zhang, L. Liu, S. Wang, Y. Wang, ACS Applied Materials & Interfaces 2017, 9, 31657; b)E. Yasun, C. Li, I. Barut, D. Janvier, L. Qiu, C. Cui, W. Tan, Nanoscale 2015, 7, 10240.

[24] R. W. Woody, A. K. Dunker, Circular dichroism and the conformational analysis of biomolecules 1996, 109.

[25] a)G. Holzwarth, P. Doty, Journal of the American Chemical Society 1965, 87, 218; b)N. J. Greenfield, G. D. Fasman, Biochemistry 1969, 8, 4108; c)S. Y. Venyaminov, I. Baikalov, Z. M. Shen, C.-S. C. Wu, J. T. Yang, Analytical biochemistry 1993, 214, 17.

[26] A. Micsonai, F. Wien, L. Kernya, Y.-H. Lee, Y. Goto, M. Réfrégiers, J. Kardos, Proceedings of the National Academy of Sciences 2015, 112, E3095.

[27] A. Bujacz, Acta Crystallogr. D Biol. Crystallogr. 2012, 68, 1278.

[28] K. Liang, R. Ricco, C. M. Doherty, M. J. Styles, S. Bell, N. Kirby, S. Mudie, D. Haylock, A. J. Hill, C. J. Doonan, Nature communications 2015, 6, 7240.

[29] L. Qi, J. Lei, Y. Zhou, Q. Gao, B. Zhang, W. Lou, Z. Luo, Chemical Engineering Journal 2023, 452, 139305.

[30] a)A. Küchler, M. Yoshimoto, S. Luginbühl, F. Mavelli, P. Walde, Nature nanotechnology 2016, 11, 409; b)K. Gupta, J. A. Donlan, J. T. Hopper, P. Uzdavinys, M. Landreh, W. B. Struwe, D. Drew, A. J. Baldwin, P. J. Stansfeld, C. V. Robinson, Nature 2017, 541, 421.

[31] a)T. H. Bayburt, J. W. Carlson, S. G. Sligar, Journal of structural biology 1998, 123, 37; b)T. H. Bayburt, Y. V. Grinkova, S. G. Sligar, Nano letters 2002, 2, 853; c)T. H. Bayburt, J. W. Carlson, S. G. Sligar, Langmuir 2000, 16, 5993; d)I. G. Denisov, S. G. Sligar, Current Opinion in Structural Biology 2024, 87, 102844.

[32] a)R. Ravanfar, Y. Sheng, H. B. Gray, J. R. Winkler, Proceedings of the National Academy of Sciences USA 2023, 120 (50), e2317372120; b)R. Ravanfar, Y. Sheng, H. B. Gray, J. R. Winkler, FEBS Lett 2023, 597 (1), 59–64.

[33] M. Babu, J. Vlasblom, S. Pu, X. Guo, C. Graham, B. D. Bean, H. E. Burston, F. J. Vizeacoumar, J. Snider, S. Phanse, Nature 2012, 489, 585.

