## Supporting Information PDF for "Mechanistic Understanding of Protein–MOF Integration through Surfactant-Driven Interfacial Design"

### MATERIALS AND METHODS

#### Materials

Bovine Serum Albumin (BSA) was purchased from Fisher Bioreagent, NJ, USA. Zinc nitrate 6-Hydrate was purchased from Ward's Science, ON, Canada. Hexadecyltrimethylammonium bromide (CTAB) and 2-methylimidazole were obtained from Sigma-Aldrich, MO, USA. Glycerol monooleate (GMO) was purchased from Spectrum Chemical MFG Corp, Gardena, CA, USA. Soy lecithin was obtained from Velona Incorporate, IL, USA. Triton X-100 was obtained from ThermoFischer Scientific, MA, USA. Bradford 1X dye reagent was purchased from Bio-Rad, California, USA. All the chemicals were used as received without further purification.

#### Experimental Methods

#### Synthesis of protein@MOF and MOF

Protein@MOFs and MOFs were synthesized using previously reported coprecipitation methods,[1] at room-temperature in aqueous-phase with a $Zn^{2+}$: 2-methylimidazole (HmIM) molar ratio of 1:60. In a typical procedure, aqueous solutions of zinc nitrate hexahydrate ($Zn(NO_3)_2 \cdot 6H_2O$) and HmIM were mixed to initiate crystallization of MOF. For protein@MOFs, 10 μM Bovine Serum Albumin (BSA) as a model protein was added to HmIM, followed by the addition of zinc nitrate to initiate encapsulation. The final mixture maintained a $Zn^{2+}$:HmIM ratio of 1:60. Surfactants, including GMO, CTAB, lecithin, and Triton X-100, were introduced prior to MOF mineralization to modulate protein interfacial properties to a final surfactant concentration of 70 μM. The crystals were collected by centrifugation and washed three times with Milli-Q water.

**Encapsulation Efficiency using Bradford Assay**

Dried sample (1 mg) was dispersed in 5 μL water, following by the addition of 250 μL Bradford reagent was added according to the Bio-Rad protocol. The absorbance at 595 nm was used to calculate the encapsulation efficiency. The standard curve was created according to microplate standard assay from the Bio-Rad for Bradford assay for the range of 125-2000 μg/ml.

**Scanning Electron Microscopy and Elemental Analysis**

Surface morphology was investigated by scanning Electron Microscopy (SEM) using a Zeiss crossbeam 540 FIB-SEM equipped with Oxford energy-dispersive X-ray spectroscopy (EDS) system from ZEISS, 07745 Jena, Germany for elemental analysis. The SEM images were collected at 5 kV accelerating voltage and 5.9 mm working distance with side mount secondary electron detector. Samples were resuspended in water and dropped onto aluminum sample stubs and were observed after drying.

**Powder X-ray diffraction (PXRD)**

Crystalline structure of the protein@MOF was investigated using PXRD. Samples were dried using a Savant Speed Vac Plus SC110A from Savant Instruments Inc. Farmingdale, New York, USA. The samples were then placed on a sample disk and PXRD patterns were recorded at a 5°/min scanning speed and 5−60° diffraction angle by a Rigaku MiniFlex II Powder X-Ray Diffractometer with Cu Ka radiation set at 30 kV and 15 mA from Rigaku Americas Corporation, 9009 New Trails Drive, The Woodlands, TX, USA.

**Thermal Gravimetric Analysis (TGA)**

Thermal stability of the samples was investigated using TGA. The samples (10 mg) were placed on a sample pan and heated in a Nitrogen atmosphere from 25 °C to 800 °C at a rate of 10 °C/min using a Shimadzu DTG-60H simultaneous DTA-TG apparatus from Shimadzu Corporation, Kyoto, Japan. The char yield was calculated according to the weight loss (%).

**Growth Kinetics Measurements**

Time-resolved size growth of the protein@MOF composites was investigated by dynamic light scattering (DLS) mode of Malvern Zetasizer pro from Malvern Panalytical Ltd., Grovewood Road, Malvern, Worcestershire, WR14 1XZ, United Kingdom. This study was performed using disposable plastic cuvettes provided by Malvern. The study was carried out through synthesis of protein@MOF in the presence and absence of surfactants with a final protein concentration of 1 μM. Data collection initiated immediately after mixing all components. Measurements were conducted for 10 cycles with optimized instrument settings, positioning at the center, attenuation of 5, 15 runs, and run duration of 0.86 seconds. All surfactant solutions were prepared in 50% ethanol. Experiments were done in triplicates. The size growth was recorded for 300 seconds, and the growth was fitted to the exponential equation using MATLAB to calculate the initial rates for the protein@MOF in the presence and absence of different surfactants.

**Zeta (ζ) Potential Analysis**

Zeta potential analysis was conducted using the static light scattering (SLS) mode of Malvern Zetasizer Pro using folded capillary zeta cell (DTS1070) cuvettes supplied by Malvern. The final concentration of protein in this study was 10 μM.

**Circular Dichroism Spectroscopy (CD)**

Secondary structure of 1 μM BSA in the presence of surfactant solutions was investigated using a J-815 CD, Tokyo, Japan. Secondary structure calculations were carried out with BeStSel.[2] The raw ellipticity data was in millidegrees with the path length of 0.1 cm. Single spectrum calculation was conducted on the data on 195-250 nm wavelength range with a scale factor of 2.

**Attenuated total reflection Fourier transformed infrared spectroscopy (ATR-FTIR)**

ATR-FTIR was performed on a Nicolet iS10 FT-IR with a Smart iTX ATR sampling accessory from ThermoFisher Scientific, Thermo Electron Scientific Instruments LLC, Madison, WI, USA.

**Ultraviolet–visible (UV-vis) Spectrophotometry**

UV-vis spectrophotometry was conducted in the range of 190-900 nm, with 1.0 nm data interval, and 0.004 s averaging time on an Agilent Cary UV-Vis Compact Peltier from Agilent Technologies, Mulgrave, Australia.

**Brunauer–Emmett–Teller (BET) Analysis**

Surface area analysis was performed using $N_2$ adsorption-desorption isotherms at 77K with a Quantachrome® ASiQwin™ analyzer. Prior to measurements, samples were degassed under vacuum at 100°C for 12h to eliminate adsorbed gases and moisture. The specific surface area of the samples was determined using the Brunauer–Emmett–Teller (BET) method. The BET equation was applied in the relative pressure range ($P/P_0$) of 0.05–0.50, where the isotherms showed linearity. The surface area was calculated based on the measured adsorption data. Pore size distribution and pore volume were further analyzed using Density Functional Theory (DFT) applied to the $N_2$ adsorption-desorption isotherms. The analysis was carried out using $N_2$ at 77 K on carbon (cylindrical pores, QSDFT adsorption branch). Model selection was based on the expected pore shape determined from SEM.

**Enzyme Activity Assay**

HRP@MOF was resuspended using brief sonication followed by vortexing to ensure uniform dispersion. To initiate the enzymatic reaction, hydrogen peroxide ($H_2O_2$) and o-phenylenediamine (OPD) as substrate were added. After incubation at room temperature for 15 minutes, the absorbance of the oxidized product was measured at 450 nm to assess the peroxidase activity of the encapsulated enzyme.

**Computational Methods**

Molecular dynamics (MD) simulations were carried out to investigate the interactions between BSA and GMO. These simulations helped identify the protein residues that interact most strongly with the surfactant molecules. The structure of BSA was obtained from the Protein Data Bank (PDB ID: 4F5S). BSA was placed at the center of a cubic simulation box (117 × 117 × 117 $Å^3$), and seven GMO molecules were randomly placed around the protein using PACKMOL software,[3] ensuring that all atoms of each GMO maintained a minimum distance of 5 Å from every protein atom. Periodic boundary conditions were applied in all directions throughout the simulations. To mitigate the sensitivity of the simulation results on the initial positions of the surfactant, six different simulation systems were prepared, each with a distinct arrangement of GMO molecules around the BSA. Additionally, a separate simulation without any GMO molecules was performed to examine the dynamical behavior of BSA in the absence of surfactants. $Na^+$ ions were added to neutralize the net charge of the systems. In a separate series of simulations (six in total), a 0.15 M concentration of NaCl was added to test the effects of ion concentration on protein–surfactant interactions. Additionally, to investigate the effect of

GMO concentration on its interactions with BSA, an additional simulation setup was constructed containing 100 GMO molecules in the presence of 0.15 M NaCl. The CHARMM-GUI website interface[4] was utilized to solvate the BSA-GMO complex with water molecules and ions and generate the input files for the simulations.

First, all systems were relaxed through geometry optimization. Then, a 250 ps simulation was performed in the constant NVT ensemble at 300 K temperature. The temperature of the simulation setups was controlled using the Langevin thermostat with a friction coefficient of 1 $ps^{-1}$. In the next step, for each system, a 10 ns MD simulation was conducted in the constant NPT ensemble at 300 K temperature and 1 atm pressure, which was treated as the equilibration simulation. Then, each system was subject to a 300 ns equilibration in the same constant NPT ensemble, which was treated as the production simulation. A 2 fs timestep was employed during the NPT simulations, with constraints applied to the lengths of all covalent bonds involving hydrogen atoms. The snapshots were saved in 0.1 ns time intervals. The system pressure was controlled by the Langevin piston Nose–Hoover method.[5] MD simulations were conducted using the NAMD software package.[6] The protein and surfactant molecules were treated with the CHARMM36m force field.[7] Water molecules were described using the TIP3P model.[8] Electrostatic interactions were treated with the particle mesh Ewald method,[9] and a 12 Å cutoff was applied for van der Waals interactions. The solvent accessible surface area (SASA) values of the protein in the presence and absence of the surfactant molecules were calculated using VMD 1.9.3.[10] Furthermore, to determine the spatial distribution of the GMOs' head groups around the protein, we constructed occupancy maps averaged over all six independent MD simulations without NaCl. These maps were generated based on the center of mass (COM) positions of all surfactant head groups throughout all MD trajectories, after aligning the protein backbone in each frame to the initial one as the reference structure. The analysis was carried out using VMD,[10] and the data were rendered as a three-dimensional isosurface to visualize the spatial distribution of the surfactant head groups with minimum occupancy 0.005.

To identify the protein residues with strong binding interactions with GMOs, we calculated the conformational factor for each residue ($P_i$, with $i$ being the residue number) that reflects how often each residue comes into contact with the surfactant's head group, since these interactions are more likely to occur at binding sites. The following method was employed to calculate the $P_i$'s: whenever any heavy atoms of the head group of a GMO molecule came within 5 Å of any heavy atom of residue $i$, it was counted as a contact.[11] For each residue, the total number of these contacts was recorded throughout all saved frames in all trajectories. Then, we calculated the average number of contacts per residue using the formula $\langle n \rangle = (\sum n_i)/N$, where $n_i$ is the number of contacts with the $i$th residue and $N$ is the total number of residues in BSA. The conformational factor $P_i$ for each residue was defined as $P_i = n_i/\langle n \rangle$, , which categorizes the strength of residue-GMO interactions: residues with $P_i \approx 1$ interact with the GMO at an average level; residues with $P_i \gg 1$ and $P_i \ll 1$ indicate higher or lower affinity with GMO than the average over all residues in the protein.

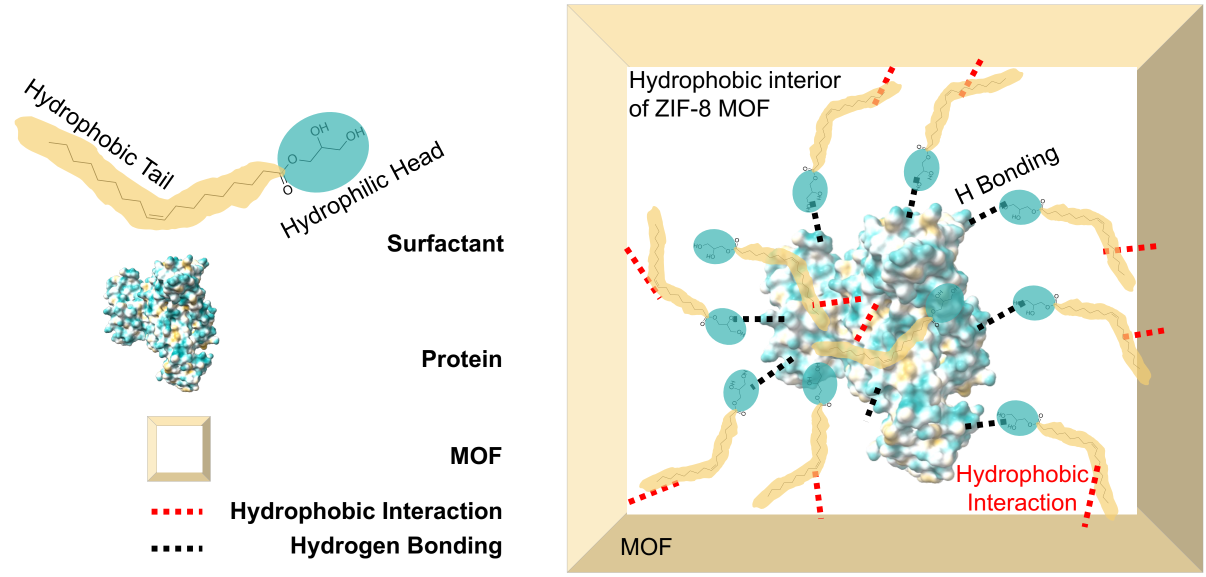

**Figure S1.** Schematic diagram illustrating specific binding interactions of surfactants at the protein–MOF interface, including potential hydrogen bonds (black dots) and hydrophobic interactions (red dots). The protein surface is colored by hydrophobicity pattern (dark cyan = most hydrophilic; dark goldenrod = most hydrophobic), using UCSF ChimeraX software, PDB ID: 4F5S. Hydrophobic interactions occur between the non-polar side chains of surface-exposed amino acids and the hydrophobic tails of surfactants. These tails may also interact with the hydrophobic interior surfaces of the ZIF-8 MOF. In parallel, the hydrophilic headgroups of surfactants can form hydrogen bonds with polar or charged amino acid residues on the protein surface, contributing to interfacial stabilization.

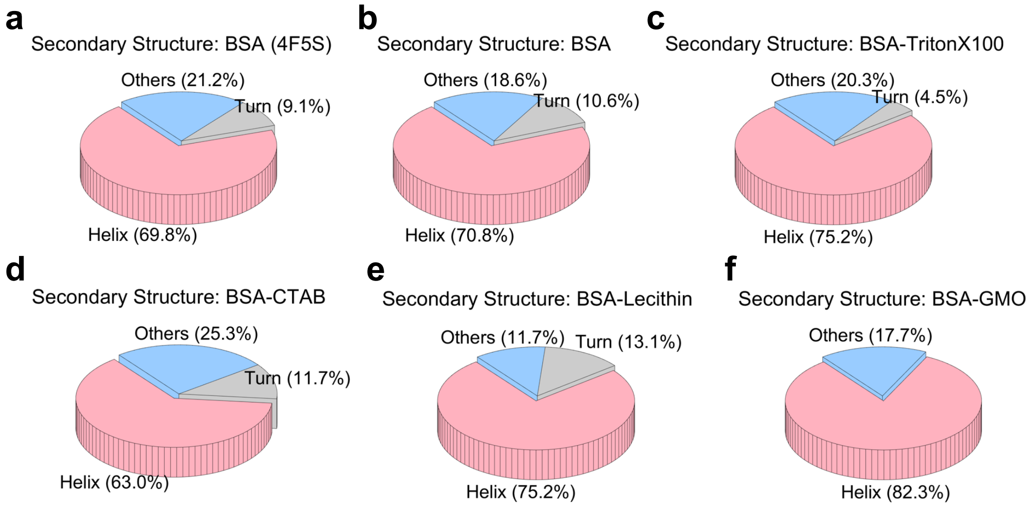


**Figure S2.** Secondary structure determination of BSA control and BSA in the presence of various surfactants.

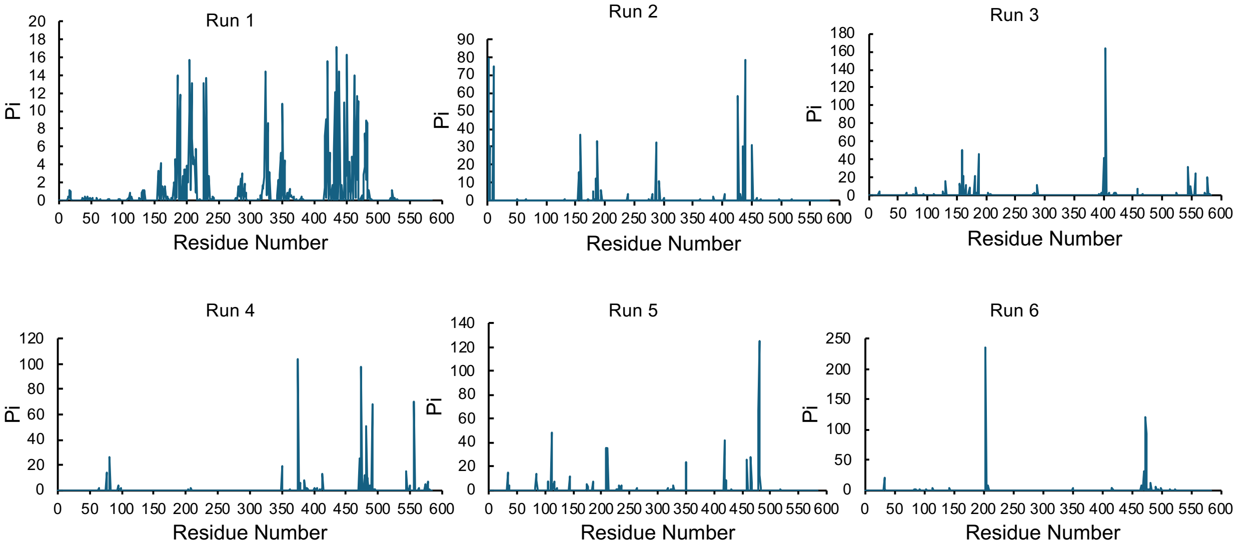


**Figure S3.** The calculated $P_i$ values for each residue of BSA across six independent MD simulations without NaCl concentration. The residues with $P_i \gg 1$ interact with the GMO molecules with above-average frequencies.

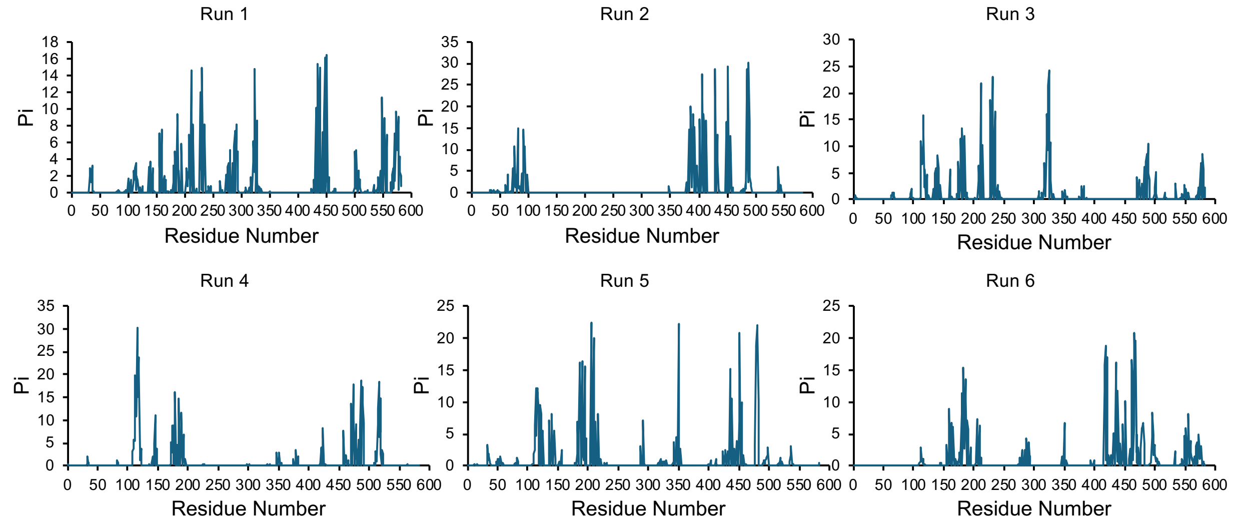


**Figure S4.** The calculated $P_i$ values for each residue of BSA across six independent MD simulations in the presence of 0.15 M NaCl. The residues with $P_i \gg 1$ interact with the GMO molecules with above-average frequencies.

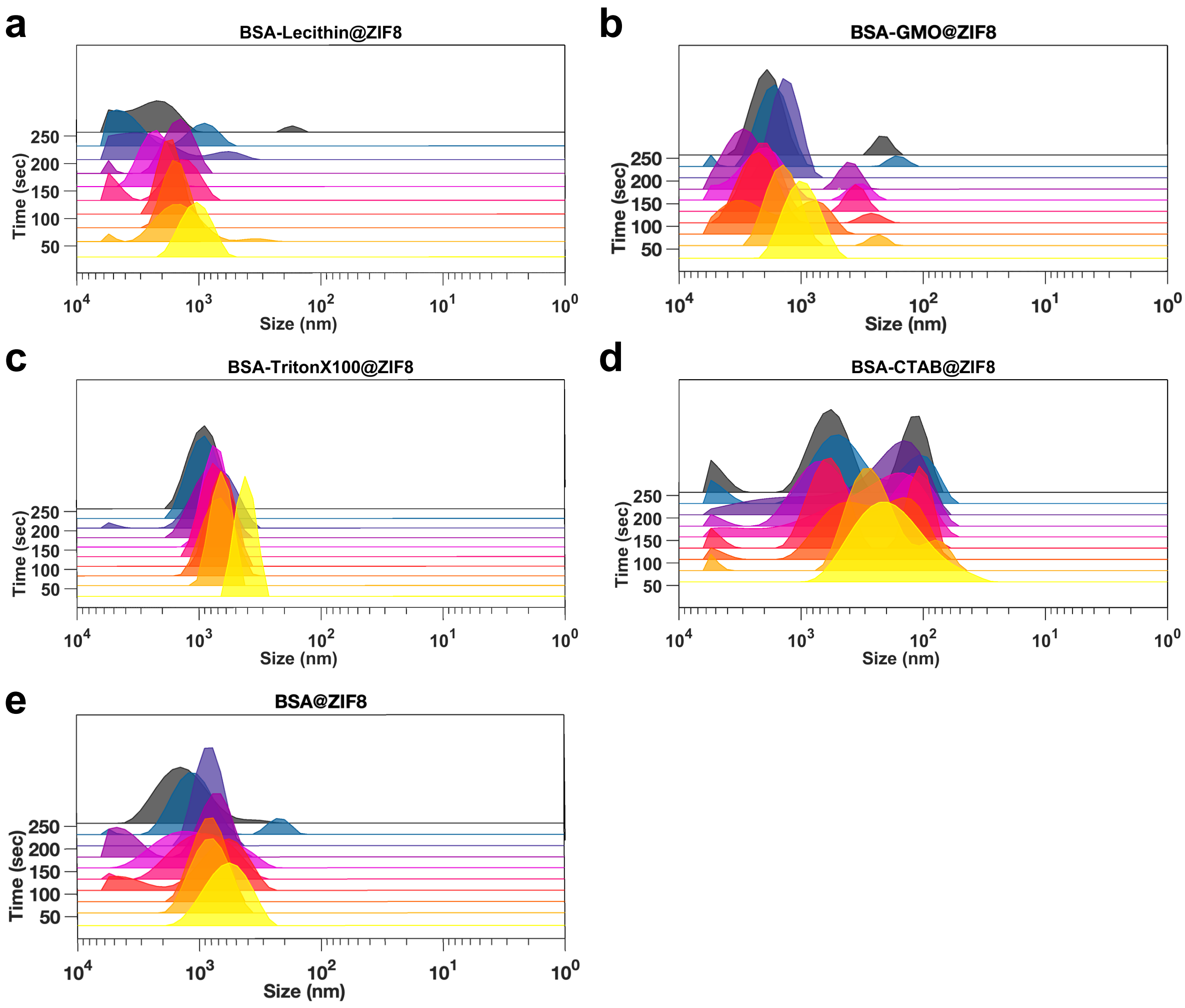


**Figure S5.** Size growth analysis using dynamic light scattering (DLS) for the formation of BSA@MOF in the presence of various surfactants in 250 seconds.

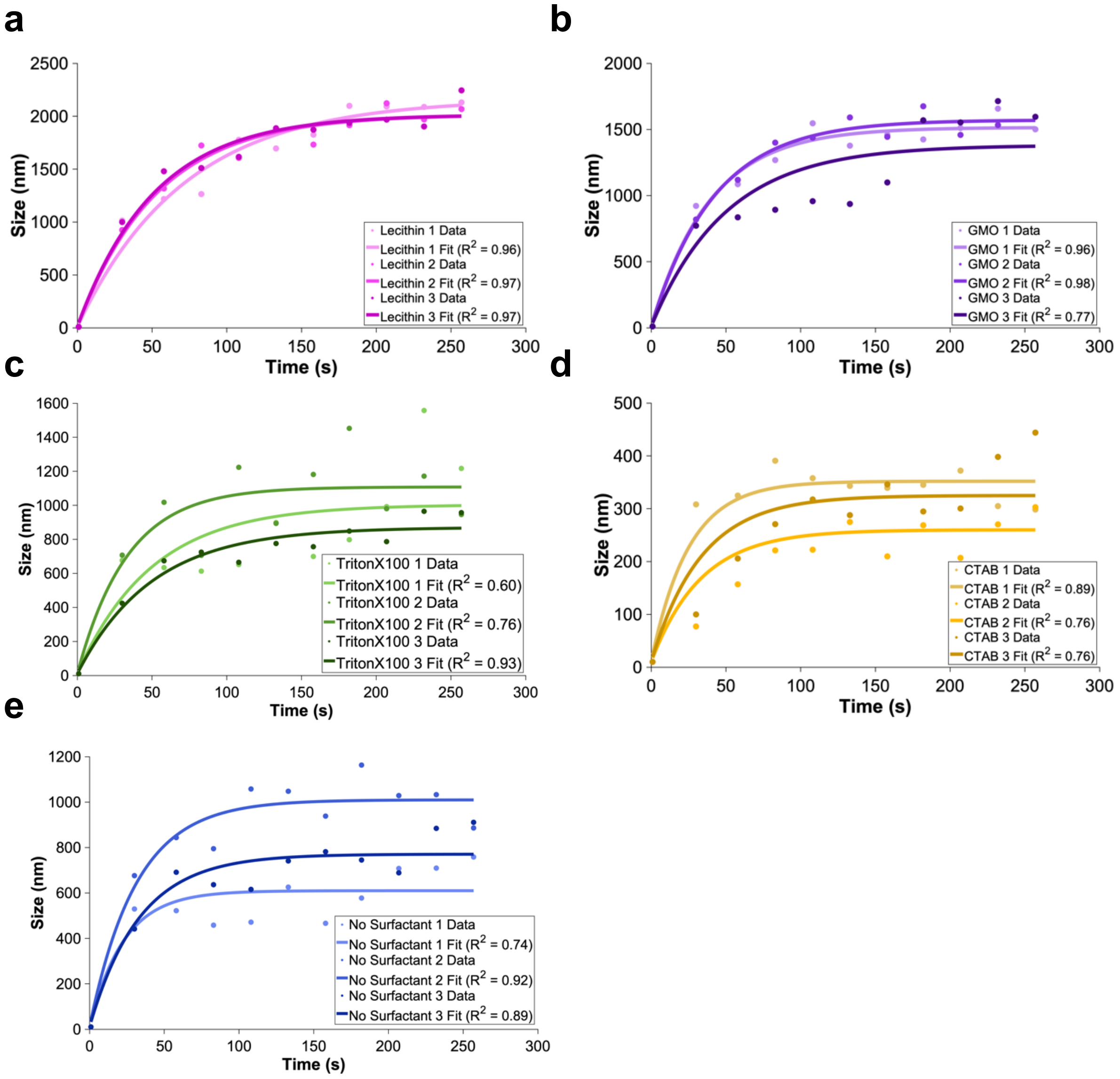


**Figure S6.** Growth kinetics with exponential fits for the formation of BSA@MOF in the presence of various surfactants to calculate initial rates.

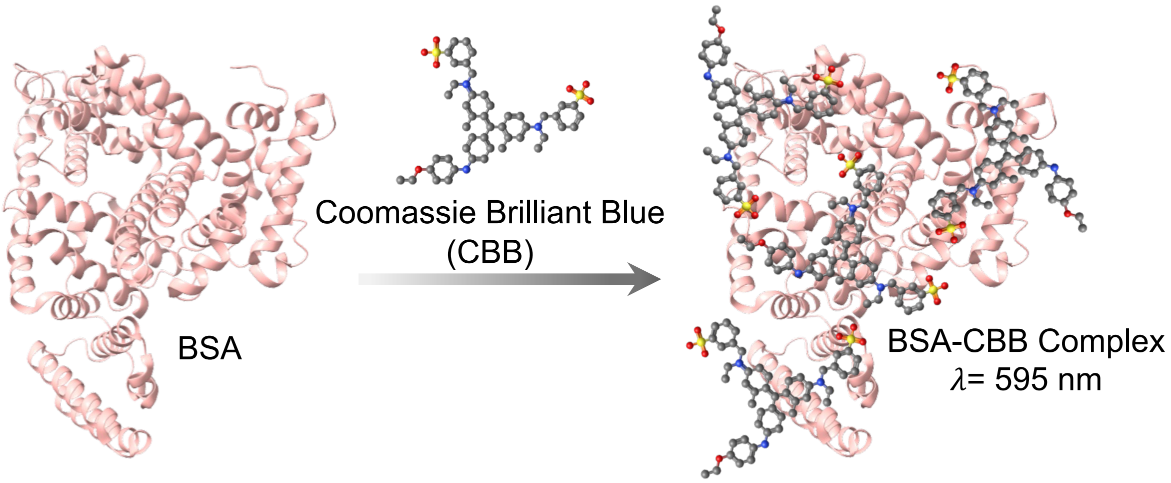

**Figure S7.** The schematic of Bradford assay, showing the interaction of Coomassie Brilliant Blue (CBB) with the protein's positive residues and formation of the colored complex with maximum absorbance at 595 nm.

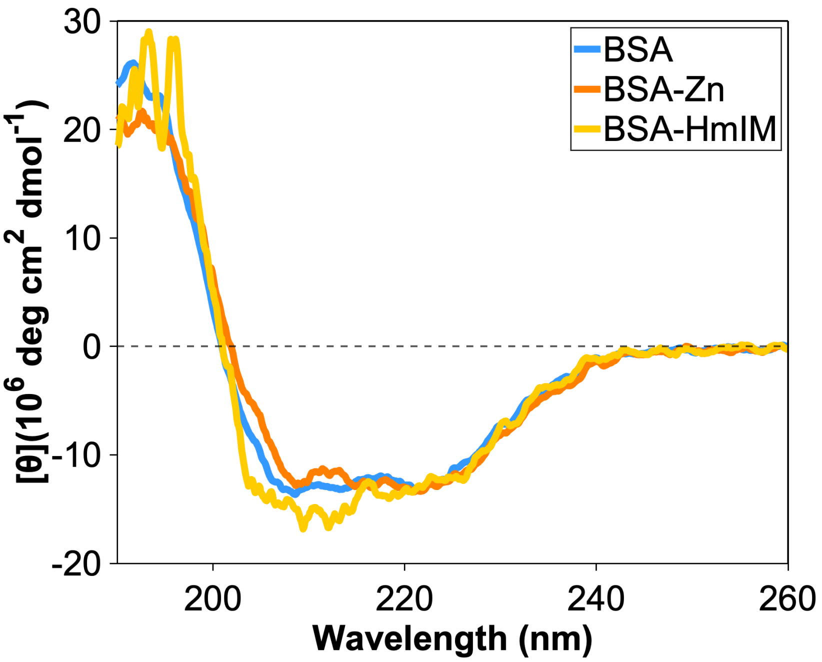


**Figure S8.** CD spectra of BSA in the presence of MOF precursors, zinc and HmIM, prior to MOF formation.

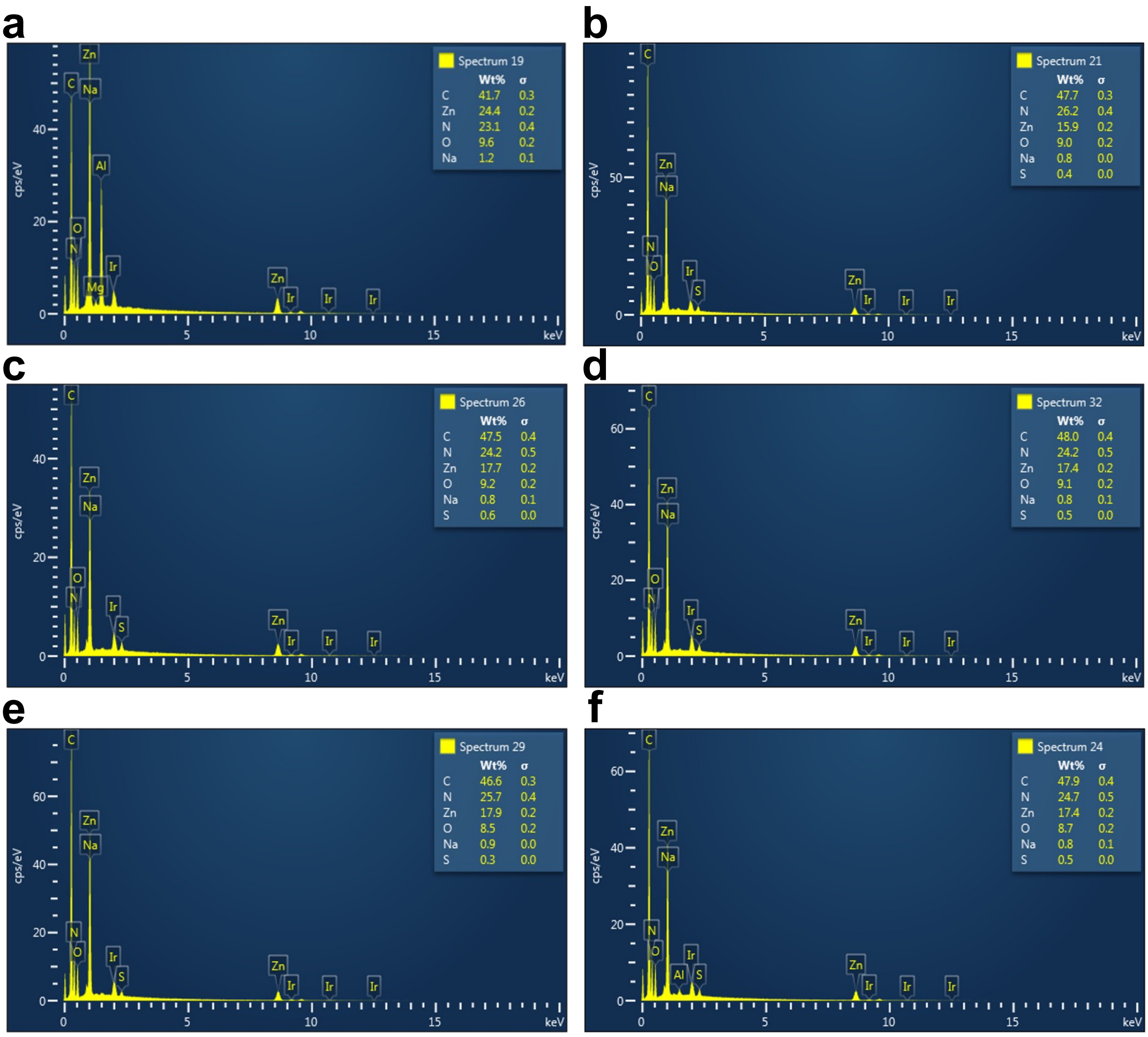


**Figure S9.** Elemental analysis of a) ZIF-8 in water, b) BSA@ZIF-8, c) BSA-GMO@ZIF-8, d) BSA-lecithin@ZIF-8, e) BSA-TritonX100@ZIF-8, and f) BSA-CTAB@ZIF-8 composites.

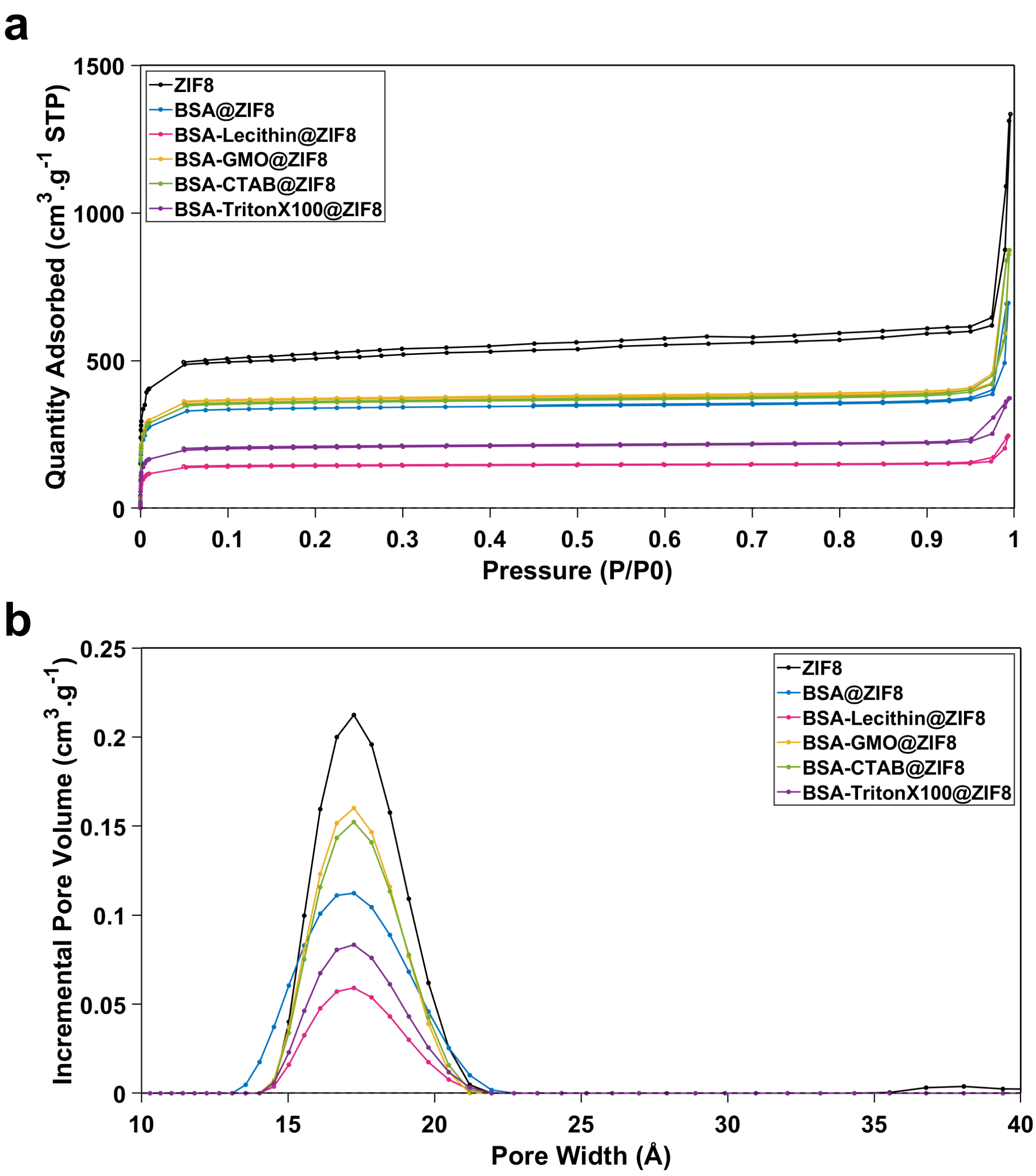


**Figure S10.** a) Nitrogen adsorption and desorption curves and (b) density functional theory (DFT) pore size distribution detected with $N_2$ adsorption and desorption at 77 K for ZIF-8 in comparison with the BSA@ZIF-8 in the presence and absence of various surfactants.

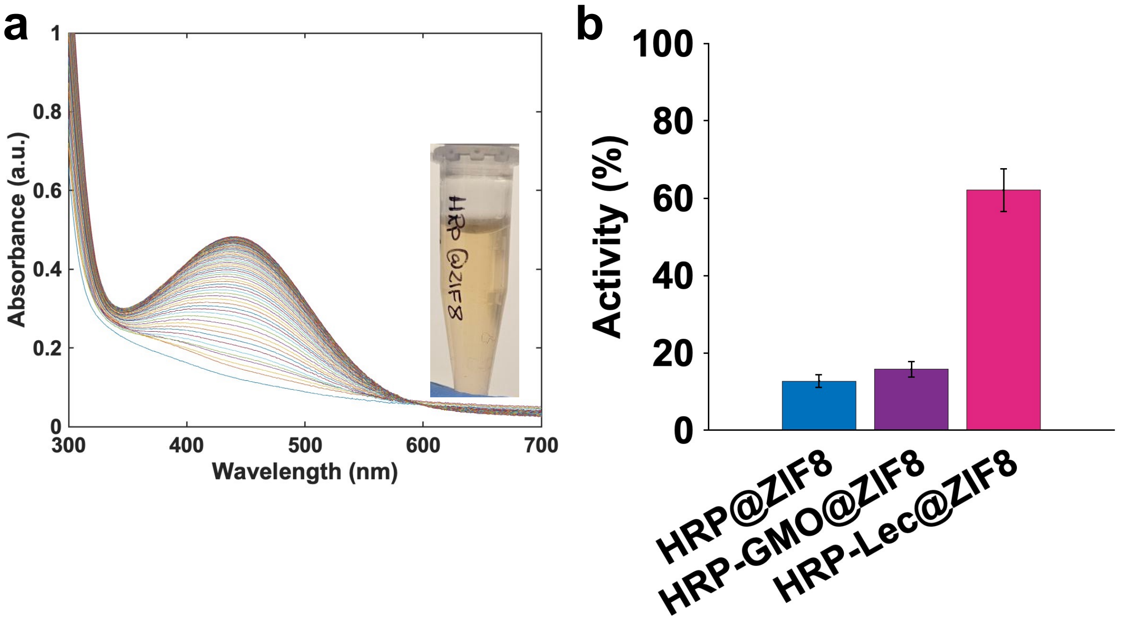


**Figure S11. a)** Representative UV-vis spectra for conversion of o-phenylenediamine dihydrochloride (OPD) to yellow oxidized product using HRP@MOF, b) HRP activity (%) improvement using lecithin and GMO.

**Table S1.** Residues with top 20 highest $P_i$ values averaged over six independent simulations with and without NaCl concentration. The residues with high $P_i$ values in both setups are highlighted in bold text.

| $P_i$ rank number | 0.15 M NaCl | | 0.00 M NaCl | |
| --- | --- | --- | --- | --- |
| | Residue | $P_i$ | Residue | $P_i$ |
| 1 | TYR451 | 9.396 | ARG435 | 16.727 |
| 2 | ARG435 | 9.16 | TYR451 | 15.885 |
| 3 | LEU115 | 7.673 | LYS439 | 15.096 |
| 4 | ALA324 | 7.581 | THR421 | 14.783 |
| 5 | THR231 | 7.478 | PHE205 | 14.708 |
| 6 | LYS439 | 7.441 | ALA324 | 13.541 |
| 7 | GLU186 | 7.285 | GLU186 | 13.169 |
| 8 | ASP323 | 7.223 | LEU462 | 13.131 |
| 9 | LYS211 | 7.197 | ARG208 | 12.904 |
| 10 | LYS116 | 6.981 | ASP323 | 12.899 |
| 11 | PRO113 | 6.661 | THR231 | 12.839 |
| 12 | LYS187 | 6.576 | PHE227 | 12.353 |
| 13 | PRO117 | 6.486 | VAL432 | 11.392 |
| 14 | LEU112 | 6.475 | LYS187 | 11.29 |
| 15 | PHE227 | 6.078 | THR466 | 11.122 |
| 16 | ALA489 | 5.982 | THR190 | 11.101 |
| 17 | GLU478 | 5.912 | LYS350 | 10.637 |
| 18 | LYS350 | 5.638 | LYS204 | 10.545 |
| 19 | PHE205 | 5.469 | VAL468 | 10.415 |
| 20 | SER479 | 5.465 | CYS447 | 10.226 |

**Table S2.** Kinetics results of protein@MOF growth in the presence of various surfactants.

| Parameters | No Surfactant | GMO | Lecithin | Triton X-100 | CTAB |
| --- | --- | --- | --- | --- | --- |

| | | | | | | |
|---|---|---|---|---|---|---|
| A (nm) | Repeat 1 | 1000.00 | 1505.16 | 2154.41 | 994.19 | 342.00 |
| | Repeat 2 | 760.89 | 1562.00 | 2005.88 | 1098.24 | 250.00 |
| | Repeat 3 | 600.00 | 1371.27 | 2005.28 | 860.77 | 315.02 |
| K (1/s) | Repeat 1 | 0.03 | 0.02 | 0.01 | 0.02 | 0.04 |
| | Repeat 2 | 0.03 | 0.02 | 0.02 | 0.03 | 0.03 |
| | Repeat 3 | 0.04 | 0.02 | 0.02 | 0.02 | 0.03 |
| AK (nm/s) | Repeat 1 | 31.97 | 38.42 | 29.97 | 19.88 | 13.68 |
| | Repeat 2 | 22.83 | 37.05 | 37.60 | 32.75 | 7.50 |
| | Repeat 3 | 26.35 | 27.42 | 38.94 | 17.22 | 9.45 |
| | Average | 27.05 ± 4.61 | 34.30 ± 5.99 | 35.50 ± 4.84 | 23.28 ± 8.31 | 10.21 ± 3.16 |

Fitting curve equations:

$S_{p@MOF} = S_0 + A(1 – e^{-kt})$

$S_0$ = 10 nm

$S_{p@MOF}$: Size of the protein@MOF (nm)

t: Time (s)

k: Size growth rate constant (1/s)

Ak: $dS_{p@MOF}/dt$ (t=0) = initial rate of growth (nm/s)

**Table S3.** Elemental analysis for MOF and protein@MOF in the presence of different surfactants.

| Surfactant | | | CTAB | Triton X-100 | GMO | Lecithin |
|---|---|---|---|---|---|---|
| Assembly | ZIF8 | BSA@ZIF8 | BSA@ZIF8 | BSA@ZIF8 | BSA@ZIF8 | BSA@ZIF8 |
| C | 41.7± 0.3 | 47.4± 0.4 | 46.5± 2.1 | 46.9± 0.2 | 47.7± 0.2 | 48.4± 1.7 |
| Zn | 24.4± 0.2 | 16.0± 0.15 | 16.9± 1.1 | 19.3± 1.2 | 17.3± 0.5 | 16.9± 0.7 |
| N | 23.1± 0.4 | 26.1± 1.2 | 25.1± 0.5 | 24.3± 1.2 | 24.4± 0.3 | 23.6± 0.7 |
| O | 9.6± 0.2 | 9.2± 1.0 | 9.6± 1.7 | 8.0± 0.5 | 9.2± 0.2 | 9.8± 0.7 |
| Na | 1.2± 0.1 | 0.8± 0 | 0.8± 0 | 0.9± 0.1 | 0.9± 0.1 | 0.8± 0.1 |
| S | - | 0.4± 0.1 | 0.5± 0 | 0.5± 0.1 | 0.5± 0.1 | 0.5± 0 |

**Table S4.** Surface area, pore width, and pore volume of ZIF-8 in comparison with protein@ZIF-8 in the presence and absence of various surfactants, derived from multipoint Brunauer–Emmett–Teller (BET) and Density Functional Theory (DFT) analysis.

| Sample Name | Surface Area ($m^2/g$) | Correlation Coefficient | Pore width (Å) | Pore Volume (cc/g) | Fitting Error (%) |
|---|---|---|---|---|---|
| ZIF8 | 2060 | 0.99 | 17.25 | 0.79 | 6.62 |
| BSA-GMO@ZIF8 | 1517 | 0.99 | 17.25 | 0.55 | 7.03 |
| BSA-CTAB@ZIF8 | 1462 | 0.99 | 17.25 | 0.54 | 6.91 |
| BSA@ZIF8 | 1384 | 0.99 | 17.25 | 0.51 | 6.85 |
| BSA-TriotonX100@ZIF8 | 829 | 0.99 | 17.25 | 0.31 | 6.71 |
| BSA-Lec@ZIF8 | 582 | 0.99 | 17.25 | 0.22 | 6.79 |

**Table S5.** Char yield (%) for free BSA, MOF, and BSA@MOF in the presence of various surfactant, extracted from the thermogravimetric analysis.

| | BSA | BSA@ZIF-8 | BSA-CTAB@ZIF8 | BSA-GMO@ZIF8 | BSA-Lecithin@ZIF8 | ZIF8 |
|---|---|---|---|---|---|---|
| Char Yield (%) | 14.8 | 33.8 | 33.0 | 37.9 | 33.6 | 36.9 |

### References


[1] a)T. Man, C. Xu, X.-Y. Liu, D. Li, C.-K. Tsung, H. Pei, Y. Wan, L. Li, *Nature Communications* **2022**, 13, 305; b)X. Wu, J. Ge, C. Yang, M. Hou, Z. Liu, *Chemical Communications* **2015**, 51, 13408; c)Y. Liu, S. Cui, W. Ma, Y. Wu, R. Xin, Y. Bai, Z. Chen, J. Xu, J. Ge, *Journal of the American Chemical Society* **2024**, 146, 12565.

[2] A. Micsonai, F. Wien, L. Kernya, Y.-H. Lee, Y. Goto, M. Réfrégiers, J. Kardos, *Proceedings of the National Academy of Sciences* **2015**, 112, E3095.

[3] L. Martínez, R. Andrade, E. G. Birgin, J. M. Martínez, *J. Comput. Chem.* **2009**, 30, 2157.

[4] S. Jo, T. Kim, V. G. Iyer, W. Im, *J. Comput. Chem.* **2008**, 29, 1859.

[5] a)G. J. Martyna, D. J. Tobias, M. L. Klein, *J. Chem. Phys.* **1994**, 101, 4177; b)S. E. Feller, Y. Zhang, R. W. Pastor, B. R. Brooks, *J. Chem. Phys.* **1995**, 103, 4613.

[6] J. C. Phillips, D. J. Hardy, J. D. C. Maia, J. E. Stone, J. V. Ribeiro, R. C. Bernardi, R. Buch, G. Fiorin, J. Hénin, W. Jiang, R. McGreevy, M. C. R. Melo, B. K. Radak, R. D. Skeel, A. Singharoy, Y. Wang, B. Roux, A. Aksimentiev, Z. Luthey-Schulten, L. V. Kalé, K. Schulten, C. Chipot, E. Tajkhorshid, *J. Chem. Phys.* **2020**, 153, 044130.

[7] a)J. Huang, S. Rauscher, G. Nawrocki, T. Ran, M. Feig, B. L. de Groot, H. Grubmüller, A. D. MacKerell, *Nat. Methods* **2017**, 14, 71; b)R. B. Best, X. Zhu, J. Shim, P. E. M. Lopes, J. Mittal, M. Feig, A. D. MacKerell, *J. Chem. Theory Comput.* **2012**, 8, 3257; c)A. D. MacKerell, Jr., M. Feig, C. L. Brooks, *J. Am. Chem. Soc.* **2004**, 126, 698; d)A. D. MacKerell, D. Bashford, M. Bellott, R. L. Dunbrack, J. D. Evanseck, M. J. Field, S. Fischer, J. Gao, H. Guo, S. Ha, D. Joseph-McCarthy, L. Kuchnir, K. Kuczera, F. T. K. Lau, C. Mattos,

S. Michnick, T. Ngo, D. T. Nguyen, B. Prodhom, W. E. Reiher, B. Roux, M. Schlenkrich, J. C. Smith, R. Stote, J. Straub, M. Watanabe, J. Wiorkiewicz-Kuczera, D. Yin, M. Karplus, *J. Phys. Chem. B* **1998**, 102, 3586; e)J. B. Klauda, R. M. Venable, J. A. Freites, J. W. O'Connor, D. J. Tobias, C. Mondragon-Ramirez, I. Vorobyov, A. D. MacKerell, R. W. Pastor, *J. Phys. Chem. B* **2010**, 114, 7830.
[8] W. Jorgensen, J. Chandrasekhar, J. Madura, R. Impey, M. Klein, *J. Chem. Phys.* **1983**, 79, 926.
[9] T. Darden, D. York, L. Pedersen, *J. Chem. Phys.* **1993**, 98, 10089.
[10] W. Humphrey, A. Dalke, K. Schulten, *J. Mol. Graph.* **1996**, 14, 33.
[11] a)M. R. Housaindokht, M. R. Bozorgmehr, M. Bahrololoom, *J. Theor. Biol.* **2008**, 254, 294; b)S. Kaviani, M. Izadyar, M. Khavani, M. R. Housaindokht, *J Mol Liq* **2020**, 317, 113933.
